# COMPASS-mediated epigenetic regulation of transcription accelerates forgetting with age

**DOI:** 10.1101/2025.10.31.685883

**Authors:** Titas Sengupta, Rachel Kaletsky, Katherine Morillo, Shiyi Zhou, Abigail Brown, Katherine Novak, Yichen Weng, Patrick Dougherty, Amayel Ka, Coleen T. Murphy

## Abstract

The molecular regulation of forgetting and its acceleration with age are not understood. Here we discovered that active gene transcription, rather than solely mRNA translation, is required for forgetting. We found that the subunits of SET1/COMPASS, a conserved histone methyltransferase complex associated with active transcription, are upregulated in aged *C. elegans* neurons, mirroring their age-related increase in mammalian brain regions governing memory. Reduction of SET1/COMPASS components blocks forgetting in both young and old animals. Optogenetic manipulation, degron-mediated conditional knockdown, pharmacological and genetic inhibition, neuronal RNA-Seq, and neuronal chromatin profiling reveal that SET1/COMPASS promotes activity-dependent *de novo* transcription in an olfactory sensory neuron that erases the associative memory trace in downstream neurons via neuropeptide signaling, resulting in forgetting. Increased SET1/COMPASS-dependent chromatin accessibility at these loci primes transcription, accelerating forgetting with age. Rescue with human SET1A and an inhibitor of human COMPASS underscore the conservation of this pathway from worms to humans. Our results identify SET1/COMPASS-mediated *de novo* gene transcription as a mechanism of forgetting, implicating increased expression of COMPASS components as a conserved driver of cognitive decline with age.

## Introduction

Forgetting is often overlooked as an essential mechanism in cognition; however, it is necessary to remove memories that are no longer useful to make room for new information. Since forgetting is adaptive, active mechanisms have evolved to regulate timely forgetting of memories^1^. That forgetting is an active process, distinct from memory maintenance pathways, has only recently been appreciated^2^, and it is critical to study the molecular regulation of active forgetting in order to understand how the durability of memories change with age. Studies in Drosophila and mice uncovered Rac1-based signaling mechanisms of forgetting^3,4^. In *C. elegans*, we and others have shown that mRNA translation is required for forgetting^5,6^ and additional studies have shown that diacylglycerol and TIR1/JNK1 signaling regulate forgetting^7–9^. However, whether epigenetic or transcriptional mechanisms are necessary for forgetting of short-term memories, whether this is the basis of a “forgetting program”, and how forgetting pathways change with age, remain unknown.

Accelerated forgetting is a hallmark of cognitive aging across species^10,11^. However, the extent to which this age-dependent memory loss is driven by active forgetting pathways, as opposed to declines in memory formation or encoding, and the molecular mechanisms that underlie increased forgetting with age, remain unclear. Defining how active forgetting is regulated at the molecular level and how these mechanisms are altered with age is critical for understanding cognitive decline and identifying targets to preserve memory function in aged individuals.

Histone post-translational modifications are significantly altered with age and in neurodegenerative diseases^12–15^ and are therefore ideal candidates for the epigenetic regulation of age-related cognitive decline. For example, histone acetylation underlies age-related impairment in synaptic plasticity and long-term memory formation^16–18^. However, whether histone posttranslational modifications regulate active forgetting, is not known.

Trimethylation of Histone 3 at lysine 4 (H3K4me3) is a conserved and abundant histone methylation mark associated with gene transcription^19,20^. In *C. elegans*, the deposition of H3K4me3 is catalyzed by a conserved SET1/COMPASS histone methyltransferase complex that includes SET-2/SET1A, WDR-5.1/WDR5, and ASH-2/ASH2L^21^. The *C. elegans* SET1/COMPASS complex acts in the germline to regulate lifespan via modulation of intestinal lipid composition^21–23^ and regulates transgenerational epigenetic inheritance of lifespan^24^. However, whether this conserved methyltransferase complex acts in the adult nervous system, and whether it contributes to age-related cognitive decline, is unknown.

Here, we discovered that active gene transcription is required for forgetting of short-term memory, raising the question of how this transcription is regulated and how it changes with age. We identified a new role for the conserved SET1/COMPASS histone methyltransferase in the regulation of short-term associative memory: our data demonstrate that SET1/COMPASS-mediated H3K4 trimethylation poises genes for transcription required during the forgetting phase of this associative memory. Dynamic changes in neural activity during memory formation and decay drive *de novo* transcription of neuronal genes, including a neuropeptidergic forgetting signal that erases the associative memory trace. Short-term memory formation and forgetting have been shown to engage translational and post-translational mechanisms^3,5^; our data demonstrate a novel role for active gene transcription in the forgetting phase of a short-term memory. We show that this mechanism drives forgetting in both young adult and old animals: increased SET1/COMPASS-mediated *de novo* transcription with age results in accelerated forgetting, while reduction of SET1/COMPASS-mediated transcription, including with a pharmacological inhibitor of human COMPASS, extends memory in both young and old animals through reduced forgetting. Mid-life inhibition of COMPASS-mediated transcription is sufficient to counter memory decline in aged animals. Thus, our findings implicate COMPASS-mediated epigenetic regulation of forgetting as a novel, conserved, and targetable mechanism of age-related memory decline.

## Results

### Active transcription drives forgetting

We previously established that learning and formation of short-term appetitive memories in *C. elegans* are transcription-independent^5,25^; however, whether forgetting requires gene transcription was unknown and seemed unlikely due to the short time course (less than 2-3 hrs). To address whether forgetting requires active transcription, we tested the effect of blocking transcription in our standard short-term associative memory paradigm^25,26^ (**Extended data Fig. 1a**). In this assay, we briefly starve worms and then expose them to food together with a neutral odorant, butanone. The conditioned worms learn to prefer butanone (t0, immediately after conditioning). Conditioned young adult wild-type worms tested after 1 hour of conditioning remember the learned association, but this memory declines significantly by 2 hours (**Extended data Fig. 1b**). To test the role of transcription in forgetting of short-term memories, we exposed worms to the transcription inhibitor actinomycin D after conditioning; with actinomycin D treatment, we observed significant memory extension (**Fig. 1a-c**). Our results therefore implicate a new role for active gene transcription in the forgetting phase of short-term associative memory, despite the short period after memory formation.

**Figure 1.**
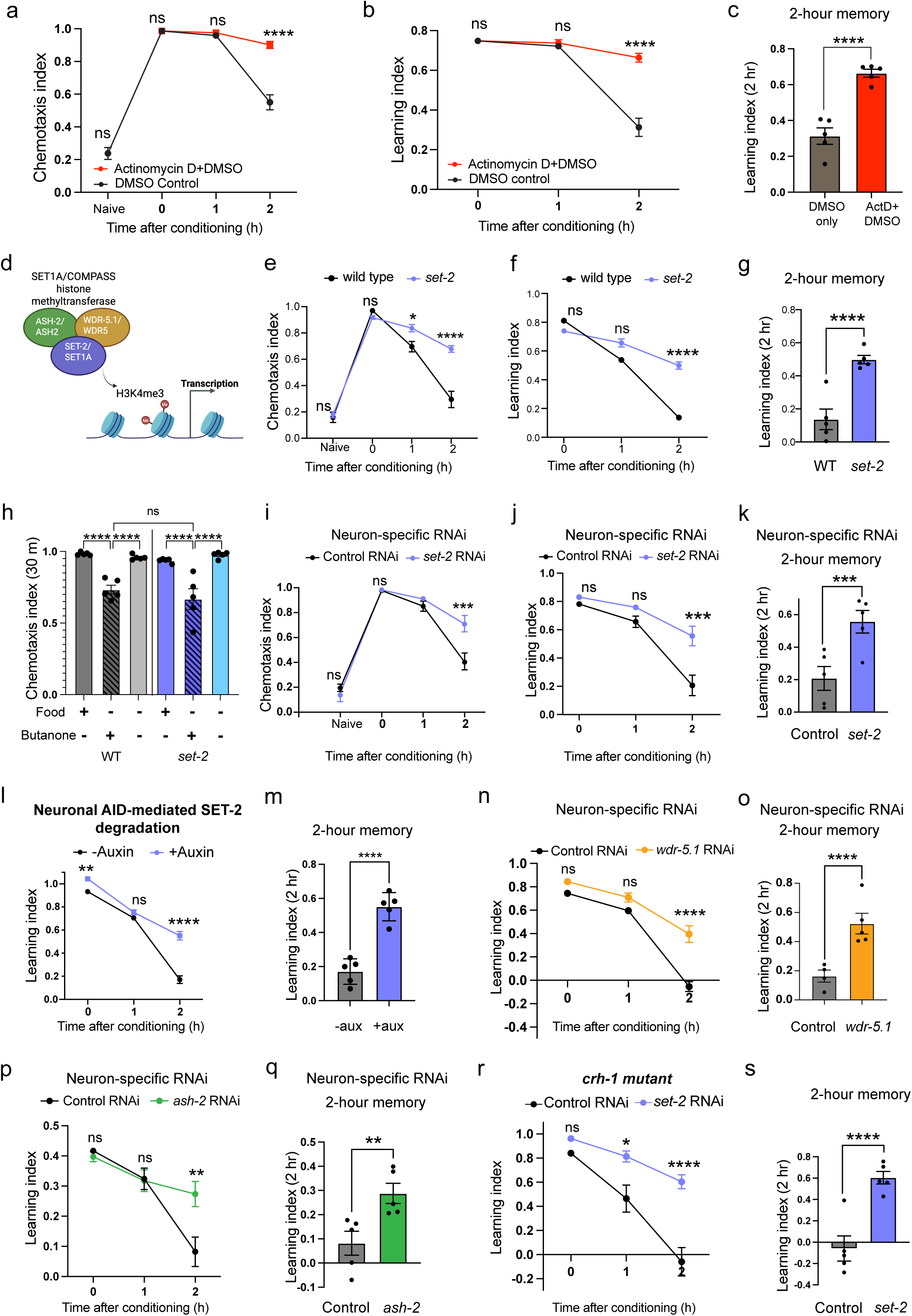
Forgetting requires new gene transcription, **(a-c)** Blocking transcription by Actinomycin D treatment (red) after conditioning (i.e., after learning) extends memory compared to solvent (DMSO) only controls (black). Chemotaxis index (a) = (# worms at butanone - # worms at ethanol)/(total worms - worms at origin). Learning index (b) = Chemotaxis index - Mean (Naive Chemotaxis index). Learning index at 2 hours after conditioning (c) represents 2-hr memory, **(d)** *set-2, ash-2,* and *wdr-5.1,* members of the SET1/COMPASS H3K4 histone methyltransferase complex, (e-g) A loss-of-function *set-2* (catalytic component of COMPASS) mutant exhibits extended memory compared to wild-type animals, **(h)** After formation of a learned food-butanone association, wild type and *set-2* mutants were (i) exposed to butanone in the absence of food (extinction training), (ii) exposed to food in the absence of butanone, or (iii) held in plates with no food or butanone (starvation control) for 30 mins. Both genotypes exhibit significant extinction of the learned preference, compared to controls, (i-k) Adult-only neuron-specific RNAi-mediated *set-2* knockdown extends memory. **(I, m)** Auxin-induced degradation (AID) of SET-2 specifically in neurons extends short-term memory, (n-q) Adult-only neuron-specific RNAi-mediated *wdr-5.1* (n, o) and *ash-2* (p, q) knockdown extend memory, **(r, s)** Neuronal RNAi-mediated *set-2* knockdown in *crh-1(n3315)* mutants also extends memory similar to wild type, (a-s) Representative experiments from three biological replicates, a, e, **i** represent chemotaxis indices, b, f, j, I, n, p, r, represent learning indices, c, g, k, m, o, q, s represent 2-hr learning indices. For 2-hr learning index plots, each dot indicates an individual chemotaxis assay plate containing -20-100 worms. 4-5 plates per time point per group were used. Two-way ANOVA with Sidák correction (a-c, e-g, i-s) or Tukey’s tests (h) was performed. Error bars = Mean±S.E.M. *P≤0.05, **P≤0.01, *\*\*\*p<* 0.001, *\*\*\*\*p*≤ 0.0001, ns - not significant.

### Adult-only, neuron-specific knockdown of *C. elegans* SET1/COMPASS components extends short-term associative memory

While learning and establishment of short-term appetitive memories in *C. elegans* are transcription-independent^5,25^, our finding that transcription is required for forgetting suggests that regulators of transcription might be involved in forgetting. The SET1/COMPASS complex is a candidate for such transcriptional regulation, as it is associated with active transcription through its trimethylation of Histone 3 at lysine 4; that is, H3K4me3 marks genes for active transcription^17^. To determine whether SET1/COMPASS plays a role in cognitive function, we tested the effect of loss-of-function mutations in *set-2*, the catalytic component of the *C. elegans* SET1/COMPASS complex, in young adult (Day 2) *C. elegans* (**Fig. 1d**). While there was no difference in learning (learning index at 0 hours) between the wild type and *set-2* mutant populations, *set-2* mutant animals have significantly better 2-hour memory of the food-odor association compared to wild-type controls (**Fig. 1e-g**). The improved memory of *set-2* mutants does not represent increased appetite, as wild type and *set-2* mutants show similar pharyngeal pumping when 2-hour memory is measured (**Extended data Fig. 1c**). Nor do *set-2* mutants have general defects in neuron function, as naïve chemotaxis to butanone (**Fig. 1e**) as well as other odorants (**Extended data Fig. 1d, e**) are unaltered. Upon extinction training, where, following the formation of a learned food-butanone positive association, worms from both genotypes were exposed to butanone without the unconditioned stimulus (food), both genotypes exhibited significant extinction of the learned preference (**Extended data Fig. 1f, g; Fig. 1h**), suggesting that the *set-2* mutants also maintain cognitive flexibility. Overall, these data suggest that *set-2* mutants exhibit extended memory without changes in general neuronal function and other aspects of cognition.

The loss-of-function *set-2* mutant has compromised SET-2 function in all tissues throughout life, which can complicate the distinction between developmental and adult effects, and effects in non-neuronal tissues vs. neurons. Therefore, we tested if adult-only knockdown (KD) of *set-2* specifically in neurons alters memory similar to the *set-2* genetic loss-of-function mutant. Worms with adult-only, neuron-specific RNAi knockdown of *set-2* have better 2-hour memory compared to controls (**Fig. 1i-k**). Similarly, conditional knockdown by auxin-induced degradation (AID) of degron-tagged SET-2 specifically in neurons extends short-term memory (**Fig. 1l, m; Extended data Fig. 2a, b**). Together, these data show that loss of SET-2 function in adult neurons extends short-term memory.

Like *set-2* knockdown, neuron-specific RNAi knockdown of two other core components of the COMPASS complex, *ash-2* and *wdr-5.1,* also improved 2-hour memory (**Fig. 1n-q; Extended data Fig. 2c, d**), suggesting that all of the COMPASS components play a role in memory extension, and that the function of the COMPASS components is necessary in adult neurons rather than in other tissues or during development for the memory-extending effect. Our data show that neuronal SET1/COMPASS plays an important role in the forgetting phase of short-term associative memory.

### CREB function is not required for SET1/COMPASS’s effects on memory

Memory is classified into different forms based on cellular and molecular requirements: short-term memory lasts for minutes to hours and requires active protein translation, while long-term memory lasts for hours (>16hrs in *C. elegans*) to days or longer in other animals and requires both gene transcription and mRNA translation into proteins^25,27^. Additionally, long-term, but not short-term, memory formation requires CREB transcription factor activity during the training period^25^. We recently discovered that *daf-2* long-lived insulin/IGF-1 signaling mutants induce neuronal CREB activity via hypodermal Notch signaling to the neurons, effectively inducing long-term, CREB-dependent memory after only a single training session^28^. To determine if the extended memory upon COMPASS knockdown similarly requires CREB function, we performed adult-only RNAi knockdown of *set-2* in *crh-1/*CREB loss-of-function mutants. However, *set-2* knockdown in *crh-1* mutants resulted in improved memory at two hours relative to controls (**Fig. 1r, s**), similar to its effect in wild-type animals (**Extended data Fig. 2e**). Therefore, CREB activity is dispensable for the extended memory exhibited upon *set-2* knockdown, consistent with *set-2* reduction extending short-term memory rather than inducing CREB-dependent long-term memory. These data indicate that *set-2* knockdown results in extended short-term, CREB-independent memory rather than a long-term, CREB-dependent memory, suggesting that SET1/COMPASS extends memory through a distinct molecular mechanism.

### SET1/COMPASS acts in distinct tissues to regulate longevity and memory

Our results demonstrate that the COMPASS complex functions specifically in neurons to regulate memory (Fig. 1). Reduction of *set-2* and other components of SET1/COMPASS in whole *C. elegans* (all tissues) or specifically in the *C. elegans* germline extends lifespan, indicating that SET1/COMPASS normally functions in the worm’s germline to limit longevity^21,23,29^. Consistent with those previous studies, we found that *set-2* knockdown in whole animals extends lifespan (**Fig. 2a**), but reduction of *set-2* only in neurons does not (**Fig. 2b**). We also found that germline-specific RNAi knockdown of *set-2* does not result in extended memory (**Fig. 2c, d**), while pan-neuronal rescue with SET-2 isoform a (SET-2a) abolished the memory improvement of *set-2* mutants **(Fig. 2e, f; Extended data Fig. 3a**). Thus, the SET1/COMPASS complex acts in neurons to regulate memory, and separately in the germline to regulate longevity; therefore, the regulation of longevity and memory by SET1/COMPASS are distinct processes.

**Figure 2.**
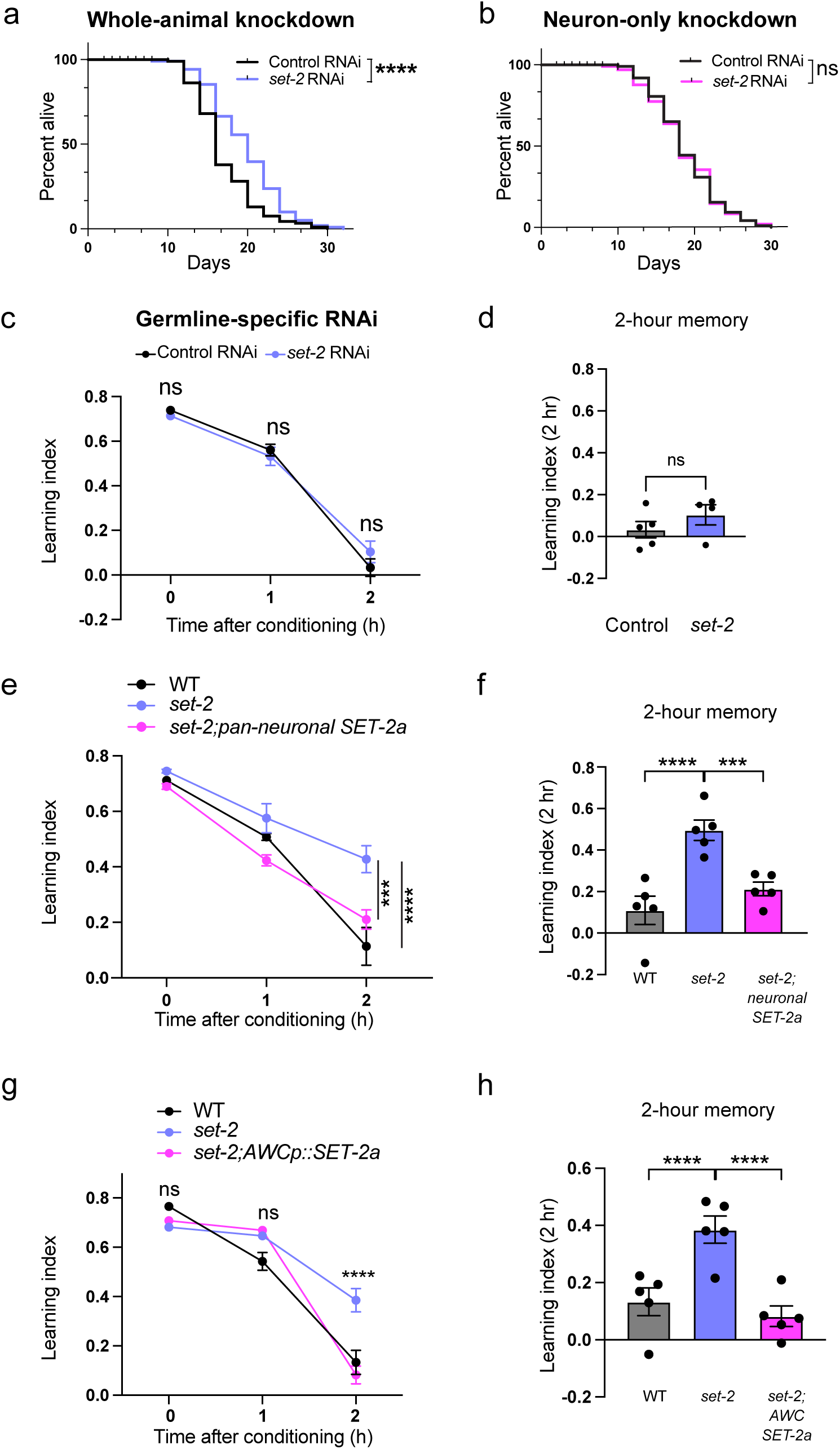
SET1 /COMPASS functions in the AWC olfactory sensory neuron to regulate forgetting, **(a, b)** RNAi-mediated knockdown of *set-2* in all tissues extends lifespan (a), while neuron-specific knockdown does not (b). **(c, d)** RNAi-mediated knockdown of *set-2* in the germline does not extend memory, **(e, f)** Pan-neuronally expressed wild-type SET-2 isoform a reverses the extended memory of *set-2* mutants, **(g, h)** Expression of wild type SET-2 isoform a in the AWC reverses the extended memory of *set-2* mutants, c, e, g represent learning indices, d, f, h represent 2-hr learning index. Two-way ANOVA was conducted and Sidák correction (for comparisons between two groups; c) or Tukey’s tests (for comparisons between more than two groups; e, g) was used for multiple comparisons, with adjusted p-values reported. Error bars = Mean±S.E.M., ***P≤0.001, ****P≤ 0.0001, ns - not significant. For the lifespan assay in a, b, *\*\*\*\*p≤* 0.0001, ns - not significant by Log-rank (Mantel-Cox) test.

### The COMPASS complex functions in the AWC neuron to regulate memory duration

We next asked in which neuron COMPASS acts to regulate memory duration and forgetting, The AWC neuron pair sense butanone and are the site of butanone-food associations^30^. Therefore, we tested if expression of the *set-2* gene specifically in the AWC sensory neuron abrogates the extended memory of *set-2* mutants. We used a neuron-specific promoter to rescue *set-2* expression, specifically in the AWC neuron pair. Similar to the pan-neuronal rescue of *set-2*, expression of SET-2a only in the AWC neurons abolished the memory improvement of *set-2* mutants (**Fig. 2f, g; Extended data Fig. 3b**). Therefore, SET-2a functions to induce forgetting of the learned association, and expression of this isoform solely in the AWC neuron is sufficient to induce this forgetting. Together, our results suggest that COMPASS-mediated active gene transcription in the AWC drives forgetting.

### The AWC neuron’s transcriptional profile

To identify *de novo* COMPASS-dependent transcriptional changes in the AWC neuron during forgetting of short-term memory, we carried out deep transcriptional profiling of the single pair of AWC neurons immediately post-training (0hr) and two hours after training (**Fig. 3a**). Because the AWC sensory neuron pair has not previously been deeply sequenced in adult animals, we first analyzed the 0 hr time point to identify basally-expressed transcripts. As expected, the transcriptome of the AWC olfactory sensory neuron is enriched in neuronal transcripts, specifically those involved in synaptic function (**Fig. 3b**). The other top functional categories of genes are protein translation and mRNA processing genes, both of which play a critical role in formation and maintenance of both short- and long-term memories^5^, and mitochondrial genes, transcription factors, DNA repair, and metabolic and cytoskeletal genes (**Supplementary Table 1; Extended data Fig. 3c**). The top most highly-expressed genes also include multiple genes encoding neuropeptide processing and maturation, dense core vesicle loading and components, and signaling machinery (**Extended data Fig. 3c**). Our deep, single-neuron transcriptomic profiling shows that the AWC, similar to mammalian neuroendocrine neurons^31^, has high expression of neuropeptide processing and release machinery.

**Figure 3.**
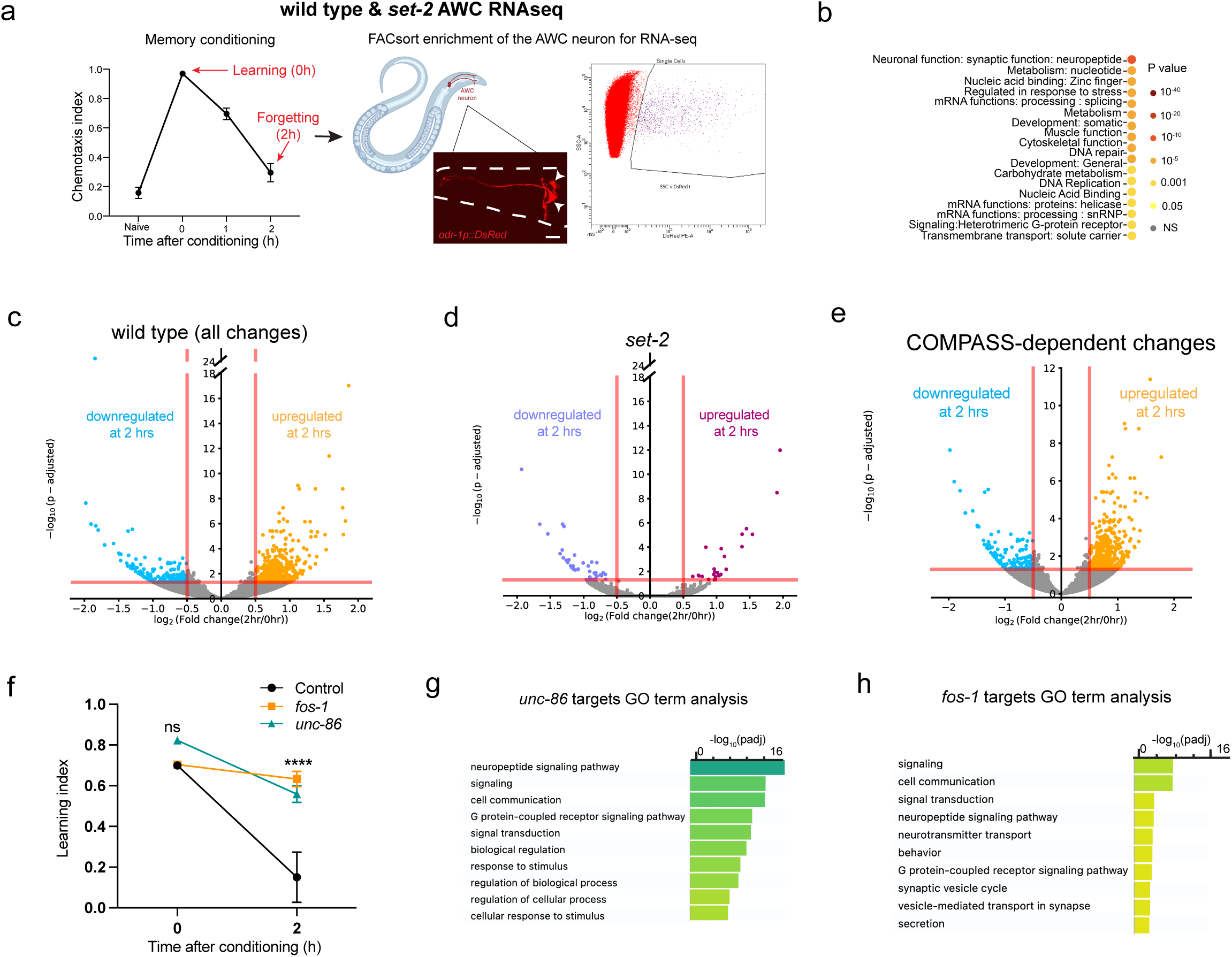
Forgetting requires SET1/COMPASS-mediated de *novo* gene transcription functions in the AWC. **(a)** AWC neurons marked with *odr-1p::DsRed* were FACS sorted and RNA-sequenced at 0 hr and 2hr post-training, (b) Functional categories of genes expressed in the AWC RNA-seq data 0-hour (after conditioning). Expressed genes were defined as those with log (Mean Normalized Counts)>2. Categories were identified using WormCAT^54^ (category 3). Top categories of genes expressed in AWC neurons are enriched for synaptic function and neuropeptides, (c-e) Differentially-expressed genes between 0 and 2 hrs after conditioning in wild type AWC (c; upregulated (orange) and downregulated (light blue)) and in *set-2* mutants (d; upregulated (purple) and downregulated (pink)) identified by DESeq2. Differentially-expressed *set-2* dependent genes (e). **(f)** *fos-1* and *unc-86* RNAi-mediated knockdown delays forgetting, **(g, h)** *unc-86* and *fos-1* targets were identified using the published motifs CATN_(3_._4)_AAAT^55^ ^56^ *(unc-86)* and TGACTCA/TTACTCA^32^ *(fos-1),* and the ‘dna-pattern’ function on RSA Tools (https://rsat.france-bioinformatique.fr/metazoa/). GO analysis was performed on *unc-86* (g) and *fos-1* (h) targets using gprofiler^57^. f represents learning indices. Two-way ANOVA was conducted and Tukey’s tests (f) were used for multiple comparisons, with adjusted p-values reported. Error bars = Mean±S.E.M. *\*\*\*\*p*≤ 0.0001, ns - not significant.

### COMPASS-dependent *de novo* transcriptional changes occur during forgetting

Next, to identify the specific COMPASS-dependent transcripts in the AWC sensory neuron pair that change during forgetting, we compared AWC gene expression profiles immediately post-training (0hr) and at two hours post-training (2hr) in wild-type (**Fig. 3c**) and *set-2* mutant animals (**Fig. 3d**). Consistent with our finding that gene transcription is required for forgetting (**Fig. 1a**), we observed that hundreds of genes are differentially expressed in the two hours between memory formation and forgetting (**Supplementary Table 2**); notably, over 95% of these gene expression changes are COMPASS-dependent, as they are unaltered in the *set-2* mutant (**Fig. 3d; Extended data Fig. 3d, e**). Since blocking transcription prevents forgetting, genes that are upregulated only in the wild-type neurons are particularly interesting, as transcription of one or more of these may drive forgetting. The COMPASS-dependent upregulated genes (**Fig. 3e**) are enriched for nuclear hormone receptors, a class of stimulus-induced transcription factors, and neuronal signaling ligands (**Extended data Fig. 3f-i**). These data suggest that an activity- and COMPASS-dependent transcriptional program alters neuronal transcription during forgetting.

### Activity-dependent transcription factors FOS-1 and UNC-86 are required for forgetting

To determine which transcription factor(s) might underlie the transcriptional changes associated with forgetting, we screened activity-associated transcription factors (TF) and TF-associated kinases for their roles in memory using neuron-specific RNAi-mediated knockdown (**Extended data Fig. 4a)**. Neuronal reduction of *fos-1* and *unc-86* extend memory (**Fig. 3f; Extended data Fig. 4b**), suggesting that these transcription factors are required for forgetting. FOS-1/FOS has conserved roles in activity-dependent transcription across organisms, and was recently shown to function as an activity-dependent TF during synaptogenesis in *C. elegans* dopaminergic neurons^32^. UNC-86/BRN3A modulates olfactory sensitivity^33^; and we previously found that *unc-86* is induced upon long-term olfactory associative memory training^27^. GO term analysis of UNC-86 and FOS-1 transcriptional target genes that are upregulated with forgetting in a COMPASS-dependent manner shows enrichment for neuropeptide signaling genes and genes encoding neuronal signaling machinery, respectively (**Fig. 3g, h; Supplementary Table 3**).

### Release of a neuropeptidergic forgetting signal actively erases the associative memory trace

Since neuronal signaling genes are upregulated during forgetting in the AWC, we next asked if release of neurotransmitters or neuropeptides from the AWC regulates forgetting. The AWC neuron uses the neurotransmitter glutamate and neuropeptides to signal to downstream neurons^34^. We first tested if glutamate release regulates forgetting by testing *eat-4* mutants, which have defective packaging of the neurotransmitter glutamate into clear core vesicles^35^. We found that *eat-4* mutants are defective for butanone associative learning (0 hr), suggesting that glutamatergic signaling from the AWC is required for learning the food-odor association (**Fig. 4a; Extended data Fig. 4c**). Since glutamatergic signaling affects learning, and learning precedes forgetting, we cannot distinguish potential roles of glutamate in forgetting. However, this is the first indication that glutamatergic signaling is required for appetitive learning in *C. elegans*.

**Figure 4.**
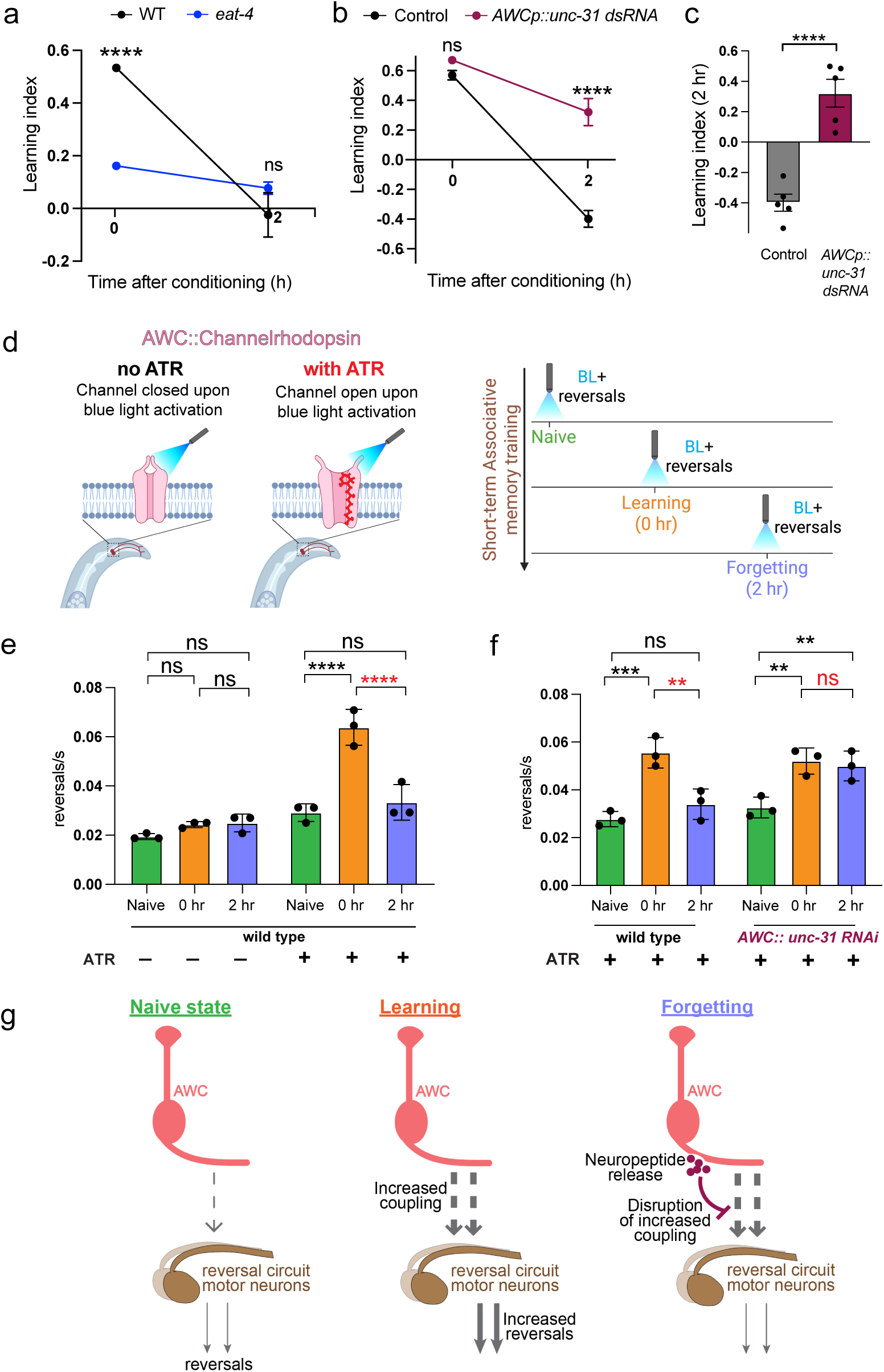
Release of neuropeptides drives forgetting by disruption of synaptic coupling. **(a)** Blocking glutamatergic release from the AWC using an *eat-4* mutant results in impaired learning **(b, c)** Blocking neuropeptide release from the AWC using a short hairpin RNA targeting *unc-31*^58^ delays forgetting **(d-g)** A strain expressing ChR2 in the AWC^37^ was activated with blue light in the naive state, after memory formation (0 h after conditioning), and during forgetting (2h after conditioning; d). Controls without all-trans retinal (ATR), where Chr2 remains in a closed conformation, show similar reversal frequencies across conditions. In the presence of ATR and blue light, reversal frequency is increased with memory formation (0 hours) and returns to basal levels during forgetting (2 hours; e, f) in wild type animals. However, upon loss of *unc-31,* no change between 0 and 2 hrs is observed in the presence of ATR (f) (g). a, b represent learning indices, and c represents 2-hr learning index. Two-way ANOVA with έidák correction (a-c) or Tukey’s multiple comparisons test (e, f) was performed, and adjusted p-values reported. Error bars = Mean±S.E.M., *\*\*P≤* 0.01, *\*\*\*p≤* 0.001, *\*\*\*\*p*≤ 0.0001, ns-not significant.

The high basal expression of multiple components of neuropeptidergic machinery in the AWC (**Extended data Fig. 3c**) may facilitate rapid processing and release of newly-transcribed peptides. To assess whether neuropeptide signaling from the AWC regulates forgetting, we reduced neuropeptide release via knockdown of *unc-31 (*which is required for all neuropeptide release^35^) specifically in the AWC; like *set-2* mutants, blocking neuropeptide release from the AWC delays forgetting, thereby extending memory (**Fig. 4b, c; Extended data Fig. 4d**). Unlike *eat-4* mutants, whole-life reduction of *unc-31* did not affect naive attraction or learning, suggesting that the neurons are largely normal, and that neuropeptide release from the AWC is specifically required for the forgetting phase.

### Forgetting disrupts learning-induced synaptic coupling between the AWC and its downstream partners

The fact that neuropeptidergic release from AWC drives forgetting suggests that remodeling of neuronal circuits may occur in downstream interneurons that are strengthened during learning. Appetitive associative learning was previously shown to increase synaptic coupling of the AWC neuron to its downstream partners in the reversal circuit^36^. This coupling was measured by optogenetic activation of the AWC in the basal state and following learning, and recording of activation-induced reversal frequencies^36^. We asked if this increased coupling is altered during forgetting. We optogenetically activated the AWC in the naïve state, upon learning (0 hr) and forgetting (2 hr), in a transgenic strain expressing Channelrhodopsin specifically in the AWC^37^ (**Fig. 4d**). Consistent with previous observations, we found that learning results in increased coupling of AWC to the reversal circuit, as measured by increased reversal frequency. We additionally found that this coupling is reduced to a near basal state upon forgetting in wild-type animals (**Fig. 4e**) but not in animals that are AWC neuropeptide release-defective (**Fig. 4f**). Taken together, our results suggest that a neuropeptidergic forgetting signal from the AWC actively erases the associative memory trace by breaking the learning-induced synaptic coupling between AWC and its downstream partners (**Fig. 4g**).

### SET1/COMPASS components are upregulated with age in the *C. elegans* nervous system

Short-term memory ability and duration decline with age in worms^25^, similar to these declines in other animals; by Day 8 of adulthood, wild-type worms have lost the ability to learn and remember, although they can still move and chemotax normally^25,26^, and by Day 5 of adulthood, worms forget the learned food-butanone association significantly faster than do young adults^25^ (**Fig. 5a-c**). This associative memory paradigm therefore captures neuronal functional changes with age, and engages evolutionarily conserved mechanisms^25,27,38^. We next asked whether members of the COMPASS complex are transcriptionally altered with age in neurons. While many neuron-specific genes decline in expression with age^38^, it is notable that the components of the COMPASS complex - *set-2/SETD1A, ash-2/ASH2L,* and *wdr-5.1/WDR5 -* are all expressed at significantly higher levels in aging neurons (**Fig. 5d**). Interestingly, the expression of *rbr-2*, which encodes the RBR-2 demethylase that counters the enzymatic activity of COMPASS by removing methyl groups from H3K4^21^, trends downwards with age (padj = 0.1169, **Fig. 5d**). The coordination of neuronal expression changes in SET1/COMPASS components with age suggests that they may play a role in increased forgetting.

**Figure 5.**
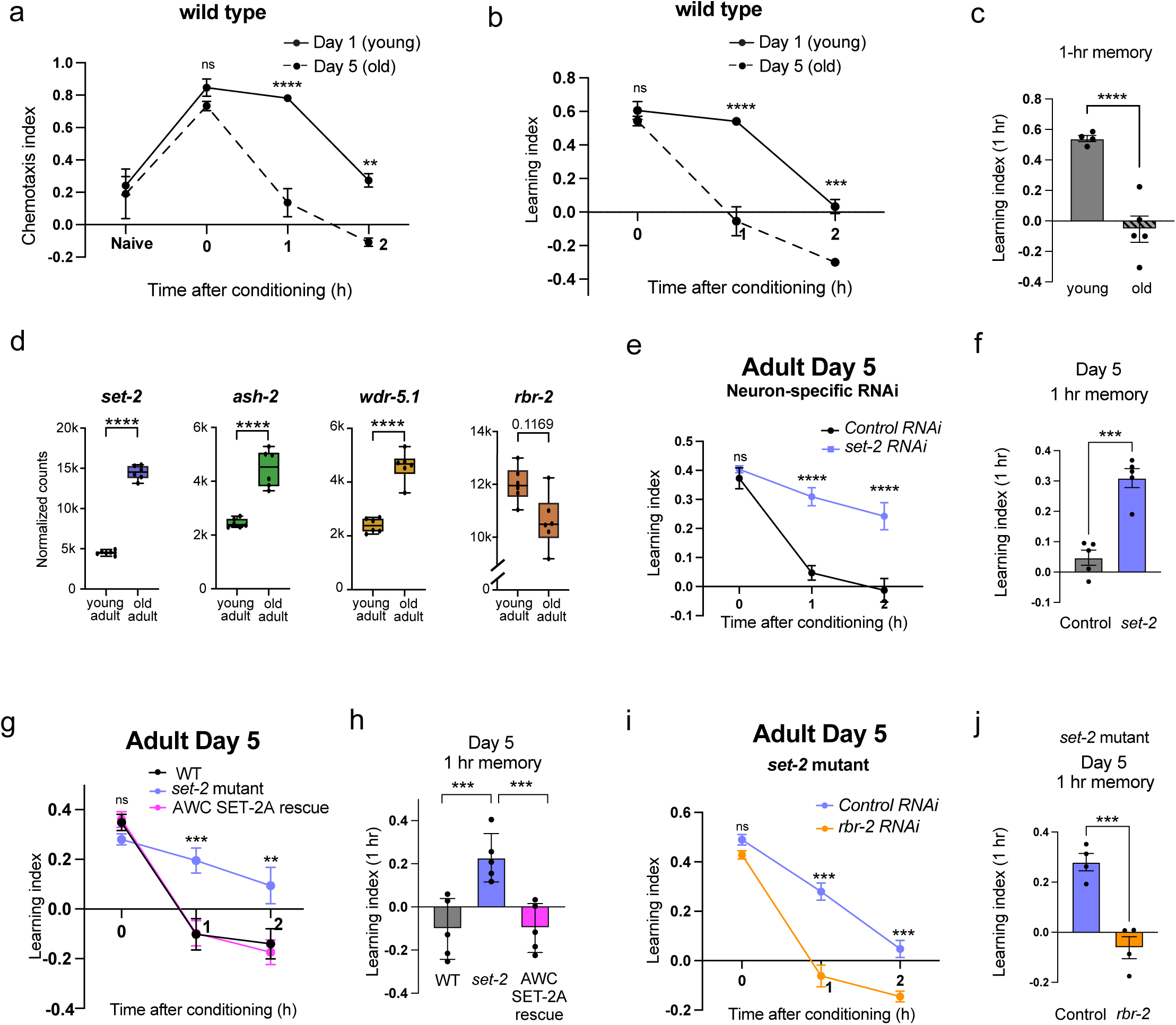
COM PASS-mediated histone methylation and transcription underlies accelerated forgetting with age. **(a-c)** Memory declines faster on Day 5 than on Day 1 of adulthood, **(d)** *set-2, ash-2,* and *wdr-5.1,* members of the SET1/COMPASS H3K4 histone methyltransferase complex, are transcriptionally upregulated in old adult (Day 8) neurons relative to young adult (Day 1) neurons, while *rbr-2,* the histone H3K4 demethylase, trends downwards (6 replicates each; padj values from DESeq2)^38^. **(e, f)** Neuron-specific RNAi knockdown of *set-2* in late larval stage extends memory in old (Day 5) adults, **(g, h)** Expression of wild-type SET-2 isoform a specifically in the AWC reverses the extended memory of *set-2* mutants in old (Day 5) adults, **(i, j)** *rbr-2* knockdown in a *set-2* mutant suppresses the extended memory of *set-2* mutants in old (Day 5) adults. A represents chemotaxis indices, b, e, g, i represent learning indices, c, f, h, j represent 1 - hr learning index. Two-way AN OVA was conducted, and Sidák correction (a, b, e, i) or Tukey’s tests (g) were used for multiple comparisons, with adjusted p-values reported. Error bars = Mean±S.E.M., **p≤ 0.01, *\*\*\*P*≤ 0.001, ****p ≤ 0.0001, ns - not significant.

### COMPASS-dependent histone methylation underlies accelerated forgetting with age

We next asked whether the COMPASS-dependent *de novo*-transcription-based mechanism of active forgetting regulates increased forgetting in aging animals; specifically, whether the faster rate of forgetting in Day 5 adults (**Fig. 5a-c**) is driven by higher expression of COMPASS components in aging animals (**Fig. 5d**). First, we tested whether reduction of *set-2* in aged adults can extend *C. elegans* short-term memory. As we observed in young animals, *set-2* knockdown in Day 5 animals robustly extends memory, while their wild-type counterparts have little to no memory at 1 hour after conditioning (**Fig. 5e, f; Extended data Fig. 5a**). The enhanced memory of *set-2* mutants in Day 5 animals is also suppressed by expression of wild-type SET-2A in the AWC (**Fig. 5g, h; Extended data Fig. 5b**). Conversely, knockdown of the *rbr-2* demethylase suppressed the enhanced memory of *set-2* mutants in Day 5 adults (**Fig. 5i, j; Extended data Fig. 5c, d**), indicating that COMPASS regulates memory through its enzymatic activity on H3K4. Together, these data indicate that COMPASS complex-mediated transcription and histone methylation regulate the forgetting of food-odor associative memory in young and aged adults, and that increased forgetting with age is at least in part due to increased levels of COMPASS components.

### Age-related neuronal chromatin changes are largely COMPASS-dependent

To query how the COMPASS complex impacts neuronal chromatin landscape changes with age, we first developed a method to assess chromatin accessibility in neurons. We performed neuronal ATAC-seq for the first time in the adult *C. elegans* nervous system by combining neuronal nuclei sorting techniques^39,40^ with adapted mammalian and whole-worm ATAC-seq methods^41,42^ (**Extended data Fig. 6a; also see Methods**). ATAC-seq profiles of young adult (Day 1) and aged (Day 8) neurons are distinct and showed significant changes in neuronal chromatin accessibility with age in wild-type animals (**Fig. 6a**), but there were fewer age-related changes in the neurons of *set-2* mutant animals (**Fig. 6b; Extended data Fig. 6b; Supplementary Table 5**). We identified about five thousand regions that are differentially accessible with age (**Fig. 6c**); by contrast, a loss-of-function mutant of *set-2* exhibits significantly fewer differentially-accessible regions (DARs) (only 1443). The vast majority (85%) of these age-related DARs are COMPASS-dependent: 2161 of the 2447 chromatin regions that show increased accessibility with age, and 2200 of the 2554 regions that show decreased accessibility with age in wild-type animals, are absent in the loss-of-function *set-2* mutant (**Fig. 6d**). Thus, SET1/COMPASS regulates the vast majority of the age-related chromatin alterations in the *C. elegans* nervous system.

**Figure 6.**
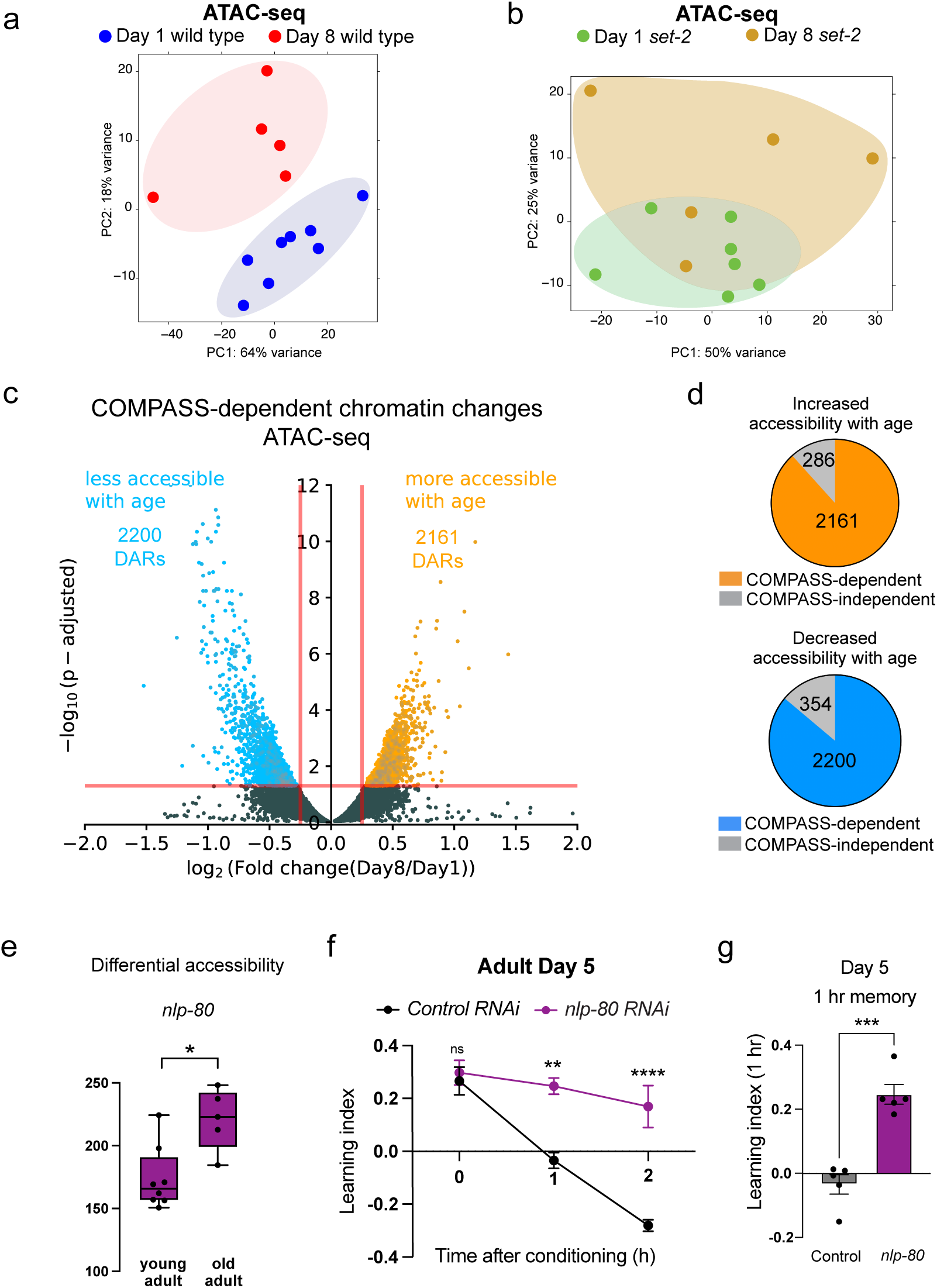
Age-related neuronal chromatin accessibility changes are largely COMPASS-dependent. **(a, b)** PCA plots for Day 1 and Day 8 neuronal ATAC-sequencing in WT (a) and *set-2* mutants (b). **(c)** Volcano plot displaying differentially accessible regions between the young adult (Day 1) and old adult (Day 8) nervous systems. The COMPASS-dependent differentially accessible regions are displayed in orange (2161; more accessible with age) and blue (2200; less accessible with age). The significantly differentially accessible regions that are COMPASS-independent are greyed out. **(d)** Proportion of regions with greater accessibility with age or lower accessibility with age that are COMPASS-independent or dependent, **(e)** Normalized read counts mapping to the peak corresponding to the *nlp-80* gene locus; p-value = padj from DeSeq2. **(f, g)** *nlp-80* is transcriptionally induced during forgetting in a COMPASS-dependent manner (Supplementary Table 2), and its reduction in aged animals extends memory (f, g). f represents learning indices, and g represents 1 -hr learning index. Two-way ANOVA was conducted, and Sidák correction was used for multiple comparisons (f, g), with adjusted p-values reported. Error bars = Mean±S.E.M., *\*\*P≤* 0.05, *\*\*P≤* 0.01, *\*\*\*p*≤ 0.001, *\*\*\*\*P* ≤ 0.0001, ns - not significant.

Using peak-to-gene mapping and a machine learning-based tissue-specific expression prediction tool^43^, we found that the gene loci that exhibit COMPASS-dependent increased accessibility with age are enriched for neuronal function (**Extended data Fig. 6c, d**). Although aging is accompanied by a general loss of neuronal identity and functions^38^, many of these neuronal gene loci exhibit increased accessibility with age in a COMPASS-dependent manner, and are therefore likely primed for transcription. These include neuronal transcription factors, membrane proteins, proteasome-related genes, and neuropeptides (including the vertebrate somatostatin-like NLP family), and a subset of these are transcriptionally upregulated upon forgetting. Such a mechanism may facilitate increased transcription from these loci with age, in turn, resulting in accelerated forgetting. Of the NLP family neuropeptides that exhibit COMPASS-dependent increased accessibility with age, *nlp-80* is transcriptionally induced during forgetting in a COMPASS-dependent manner (**Fig. 6e**; **Supplementary Table 2**). To test this model, we reduced the expression of *nlp-80* in aged animals; like *set-2* reduction, neuron-specific knockdown of *nlp-80* delays forgetting in old adults (**Fig. 6f, g; Extended data Fig. 6e**).

### Forgetting can be reversed in mid-life

Because we found that age-related changes in neuronal chromatin accessibility are driven by the activity of the COMPASS complex (whose components’ expression increases with age and drive forgetting) we next asked whether these changes could be slowed or reversed. An ideal intervention for cognitive decline would be effective when applied to older individuals to reverse or halt decline, in contrast to an intervention that must be initiated when an individual is still young. We knocked down *set-2* in neurons starting at Day 3 of adulthood and found that this mid-life treatment is sufficient to enhance memory of old Day 5 adults, suggesting that the levels of neuronal H3K4me3 can be dynamically modulated at different stages of adulthood (**Fig. 7a-c; Extended data Fig. 6f**).

**Figure 7.**
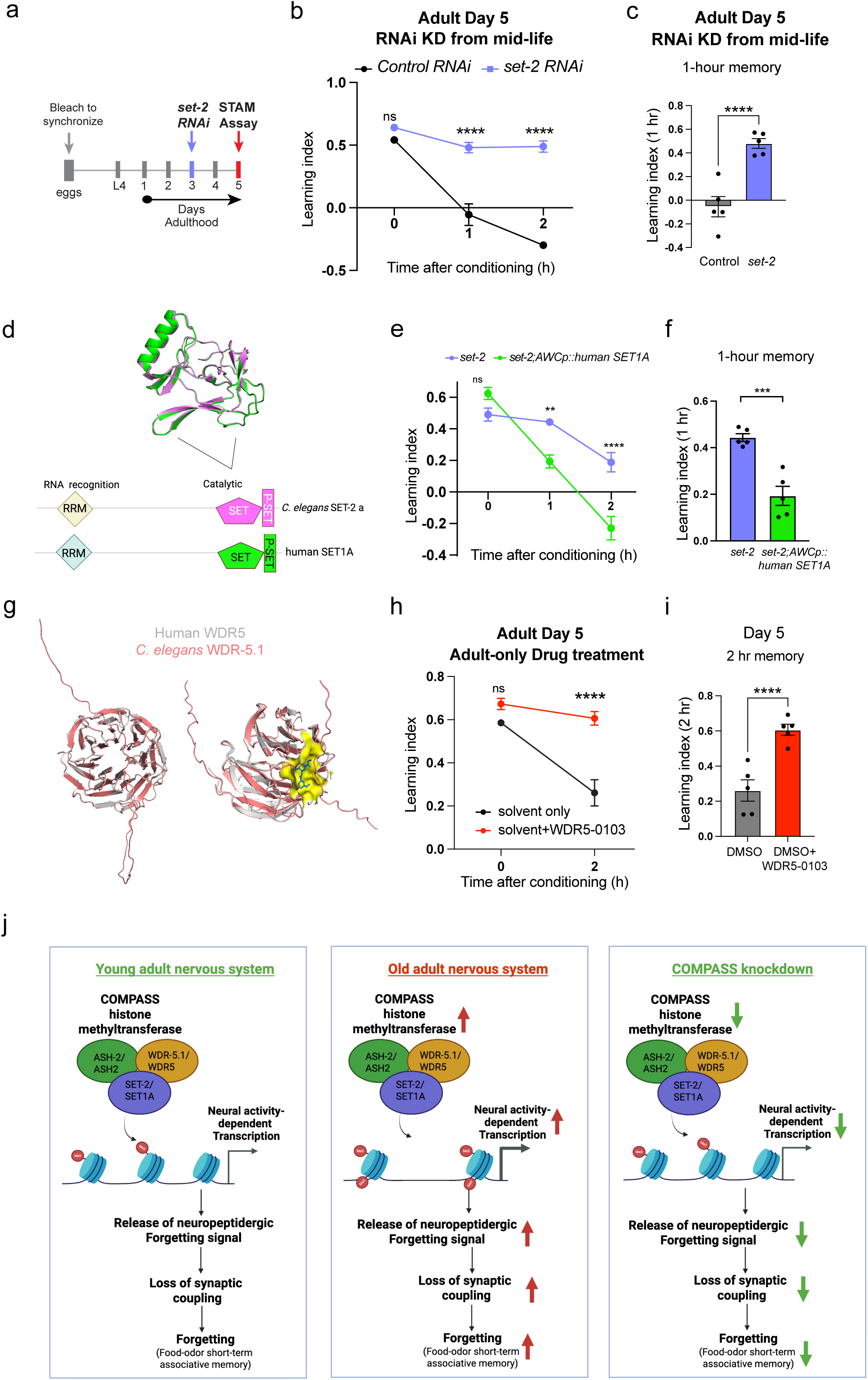
An inhibitor of human COMPASS delays forgetting and extends memory in old adults, **(a-c)** Neuron-specific RNAi knockdown of *set-2* in mid life (Day 3) extends memory in old (Day 5) adults, (d) Human SET1A, similarly to C. *elegans* SET-2 isoform a, has the SET and P-SET domains, and RNA recognition motifs (RRM). **(e, f)** Expression of Human SET1A in the AWC reverses the extended memory of *set-2* mutants, (g) Model of C. *elegans* WDR-5.1 predicts a similar binding site for the human COMPASS inhibitor, WDR5-0103. **(h, i)** Pharmacological inhibition by WDR5-0103, extends memory in old (Day 5) adults, (j) Model: COMPASS-mediated neural activity-dependent *de novo* transcription and release of neuropeptides erases the memory by disruption of synaptic coupling, thereby resulting in forgetting. In old adults, where genes transcribed during forgetting are more accessible, increased COMPASS-mediated transcription and release of neuropeptides result in accelerated forgetting. In COMPASS mutants, reduced transcription and release of neuropeptides delays forgetting, thereby extending memory, b, e, h represent learning indices, c and f represent 1 - hr learning index, and i represents 2-hr learning index. Two-way ANOVA was conducted for all the assays. Sidák correction was used for multiple comparisons, with adjusted p-values reported. Error bars = Mean±S.E.M., *\*\*P* ≤ 0.01, *\*\*\*p≤* 0.001, ****P≤ 0.0001, ns - not significant.

### Human SET1A recapitulates *C. elegans* SET-2’s role in forgetting

The SET1/COMPASS complex is very similar in composition and function across organisms, from yeast to humans^20^. SET1A, the catalytic component of the human SET1A/COMPASS methyltransferase, is the human ortholog of *C. elegans set-2; C. elegans* SET-2 isoform A closely resembles the human ortholog SET1A in sequence and domain architecture (**Fig. 7d**). To test if human SET1A and *C. elegans* SET-2 isoform a are functionally similar in our context, we expressed human SET1A in the AWC in a *set-2* mutant background. Strikingly, expression of human SET-1A in the AWC is sufficient to disrupt the extended memory of *set-2* mutant animals (**Fig. 7e, f**; **Extended data Fig. 6g**), similar to wild-type *C. elegans* SET-2 isoform a. Thus, *C. elegans* SET1/COMPASS is functionally interchangeable with human SET1/COMPASS, suggesting that the function of this complex in regulating forgetting may be deeply conserved across phyla.

### Forgetting can be blocked pharmacologically by an inhibitor of human COMPASS

We next tested whether pharmacological inhibition of COMPASS could affect memory or forgetting. The compound WDR5-0103^44^ inhibits human COMPASS function via the WDR5 subunit ortholog of worm WDR-5.1. As *C. elegans* WDR-5.1 shows near-perfect structural alignment with human WDR5 (**Fig. 7g**), and we showed that human SET1A can rescue *set-2* loss in *C. elegans* neurons, we hypothesized that this COMPASS inhibitor would also extend memory analogously to *C. elegans* COMPASS mutants. We exposed adult worms to 100 μM WDR5-0103 and tested memory on Day 5. Compared to controls, WDR5-0103-treated animals exhibit significantly improved memory (**Fig. 7h, i; Extended data Fig. 6h**), highlighting COMPASS as a druggable target that could affect cognitive aging in humans.

## Discussion

Memories are forgotten faster with age and in neurodegenerative disorders^10,45^, highlighting the importance of understanding how forgetting is regulated. Animals across phyla have evolved mechanisms that both establish and erase memories at different timescales, but why memories have evolved to last for specific lengths of time when it is molecularly feasible for them to last much longer has not been answered. If memory capacity is limited, mechanisms that remove old memories to make space for new ones would be necessary to enable responses to dynamically evolving contexts, and prevent context-irrelevant behaviors. Perhaps more importantly, mechanisms for motivated forgetting have likely evolved to allow real-time updating of information or to prevent retrieval of undesired or uninformative memories.

Much of the cognitive neuroscience field has focused on memory formation and consolidation, and there has been comparatively less emphasis on the study of forgetting itself. This is because, historically, forgetting was believed to be a passive decay of memory maintenance mechanisms, instead of being an independently regulated arm of memory. However, there is increasing appreciation for the idea that forgetting is an active process^1,46^; therefore, it is critically important to identify mechanisms that regulate active forgetting and examine how these mechanisms are altered with age to drive cognitive decline.

Short-term memories were believed to exclusively employ transcription-independent mechanisms because of their short lifetimes; however, spatially- and temporally-resolved transcriptomic profiling experiments during the course of rapid forgetting were lacking. Here, we discovered that forgetting of short-term memories requires *de novo* gene transcription on a rapid time scale (<2hrs). We found that the COMPASS complex gates this transcription, and increased COMPASS complex-mediated active transcription results in accelerated forgetting in aged animals (**Fig. 7j**). Reduced activity-dependent transcription caused by downregulation or inhibition of COMPASS components slows forgetting in young and old animals, improving memory in young adults and counteracting increased forgetting in old animals (**Fig. 7j**).

The genes induced during forgetting include conserved neural activity-responsive genes such as nuclear receptors, neuropeptides, and synaptic genes, therefore representing an activity-dependent transcriptional program that drives forgetting. Epigenetic dysregulation of this program with age accelerates forgetting. In mammals, neural activity-dependent immediate early gene (IEG) transcription occurs immediately following a training event^47^. Although short-term memories can form without IEG-mediated transcription, the gene products of activity-induced transcription can be produced rapidly enough (∼20 mins) to influence the decay of short-term memories. Therefore, although immediate early gene transcription has been primarily characterized in the context of long-term memory formation, it may play a previously undescribed role in the forgetting of short-term memories. Since gene transcription is intimately linked to neural activity and encodes neural activity patterns and histories^48,49^, a transcription-coupled mechanism may ensure context-appropriate forgetting. Transcription-based forgetting may also leave a transcription or chromatin-level footprint that competes with or primes neurons for long-term memory consolidation. We find that neuropeptides are the top functional category of the newly transcribed genes during forgetting, and blocking neuropeptide release delays forgetting. IEG-mediated transcription has been shown to produce neuropeptides across organisms^50^, and because of their short transcript lengths, these may be some of the earliest genes to be fully transcribed following activity-induced transcription induction, serving as drivers for forgetting.

At the circuit level, we observed that learning increases the coupling of the AWC olfactory sensory neuron to downstream motor circuits, and the neuropeptidergic forgetting signal reverses this coupling. This altered circuitry influences the navigation of odor gradients upon learning and forgetting. When an animal deviates from its track towards the odorant, AWC (being an odor-OFF neuron^51^) is activated, inducing reversals, therefore, bringing the animal back on the right trajectory. Learning-induced increased coupling between the AWC and the reversal circuit increases reversals upon deviation from the odor gradient, therefore allowing animals to reach the odor gradient more efficiently. Forgetting involves restoration of the increased reversal frequency back to basal levels. We identify neuropeptide signaling as a key mediator of this circuit remodeling, linking activity-dependent transcription to circuit-level and behavioral changes during forgetting.

Together, our work identifies COMPASS-mediated epigenetic priming of transcription as a conserved and reversible mechanism that accelerates active forgetting with age. SET1/COMPASS has been shown to have both enzymatic and non-enzymatic roles^52^, and levels of H3K4me3, the enzymatic product of COMPASS, increase with age in the mouse hippocampus^53^. In our context, SET1/COMPASS functions through its enzymatic activity on H3K4, as knockdown of the H3K4 demethylase, RBR-2/KDM5, suppresses the effect of COMPASS knockdown on memory. This shows that levels of a single histone modification in neurons can modulate the rate of forgetting with age. Our genetic rescue with human COMPASS and pharmacological inhibition experiments also suggest that this mechanism of forgetting may be evolutionarily well-conserved. Thus, SET1/COMPASS-dependent transcriptional regulation of forgetting may serve as a therapeutic target for the prevention of age-related short-term memory decline.

## Acknowledgments

We thank the *Caenorhabditis* Genetics Center for *C. elegans* strains. We also thank W. Wang, J. Arly Volmar, and J. Miller (Genomics Core Facility, Princeton University) and C. DeCoste and the Molecular Biology Flow Cytometry Resource Facility, which is partially supported by the Rutgers Cancer Institute of New Jersey NCI-CCSG P30CA072720-5921, for assistance; BioRender for model figure design software; Dr. Andrew Leifer and the Leifer lab (Princeton University) for optogenetics strains and advice; Dr. Navin Pokala for optogenetics advice; Madeleine Chong for assistance with experiments and analysis of memory assay data; and the Murphy lab for discussion and feedback. C.T.M. is the Director of the Lewis Sigler Institute for Integrative Genomics and the Simons Foundation for Plasticity in the Aging Brain. This work was supported by a Pioneer Award to C.T.M. (NIGMS DP1GM119167), a Simons Foundation grant to C.T.M. (AN-NC-AB-Research-00811235SPI), a Damon Runyon Fellowship to T.S., the Princeton Presidential Postdoctoral Fellowship to A.B., and T32GM148739 for support of K.M.

## Materials and Correspondence

Further information and requests for resources and reagents should be directed to and will be fulfilled by Coleen T. Murphy

## Supplementary Tables

**Supplementary Table 1**

Genes expressed (log Mean normalized counts>0) 0 hr after conditioning (AWC RNA-seq)

**Supplementary Table 2**

Differentially expressed genes during forgetting (AWC RNA-seq) (0 vs 2 hr after conditioning).

**Supplementary Table 3**

Genes upregulated during forgetting that contain *fos-1* or *unc-86* motifs.

**Supplementary Table 4**

Differentially accessible regions – Day 1 vs Day 8 neuronal ATAC-seq (wild type).

**Supplementary Table 5**

Differentially accessible regions – Day 1 vs Day 8 neuronal ATAC-seq (*set-2*).

**Supplementary Table 6**

Source data

## Methods

### General Worm maintenance

All strains were maintained following standard methods. For all experiments, worms were maintained at 20°C on plates made from nematode growth medium (NGM: 3 g/L NaCl, 2.5 g/L Bacto-peptone, 17 g/L Bacto-agar in distilled water, with 1 mL/L cholesterol (5 mg/mL in ethanol), 1 mL/L 1M CaCl2, 1 mL/L 1M MgSO4, and 25 mL/L 1M potassium phosphate buffer (pH 6.0) added to molten agar after autoclaving) or high growth medium (HGM: NGM recipe modified as follows: 20 g/L Bacto-peptone, 30 g/L Bacto-agar, and 4 mL/L cholesterol (5 mg/mL in ethanol); all other components same as NGM), with OP50 *E. coli* for *ad libitum* feeding. For auxin experiments, the standard HGM molten agar was supplemented with 2.5 mL 400mM indole-3-acetic acid (IAA): Alfa Aesar (#A10556) freshly prepared in ethanol and plates were seeded with *E. coli* for *ad libitum* feeding. Synchronized AID worms were transferred to HGM with auxin plates for 16-20 hours before behavior assay. For RNAi experiments, the standard HGM molten agar was supplemented with 1 mL/L 1M IPTG (isopropyl b-d-1-thiogalactopyranoside) and 1 mL/L 100 mg/mL carbenicillin, and plates were seeded with HT115 *E. coli* for ad libitum feeding. To synchronize experimental animals, eggs were collected from gravid hermaphrodites by exposing the animals to an alkaline-bleach solution (e.g., 1.5 ml sodium hypochlorite, 0.5 mL 5N KOH, 8.0 mL water), followed by repeated washing of collected eggs in M9 buffer ((6 g/L Na2HPO4, 3 g/L KH2PO4, 5 g/L NaCl and 1 mL/L 1M MgSO4 in distilled water). For Day 5 experiments, worms were transferred at the L4 larval stage onto HGM plates supplemented with 500ml/L 0.1M FUdR (5-Fluoro2’-deoxyuridine) for a final concentration of 0.05M FUdR and were transferred back to standard HGM 20 hours before memory assay.

### C. elegans strains

*C. elegans* strains used in this study - CQ810: *crh-1(n3315); vIs69 [pCFJ90(Pmyo-2::mCherry + Punc-119::sid-1)] V*, CQ811: *set-2 (ok952); wqEx94 [Prgef-1::set-2a+Pmyo-2::GFP]*, CQ812: *set-2 (ok952);* CQ813: *set-2 (ok952); wqEx96[Podr-1::set-2a+Pmyo-2::GFP],* LC108: *vIs69 [pCFJ90(Pmyo-2::mCherry + Punc-119::sid-1)] V,* MAH677: *sid-1(qt9) V; sqIs71 [Prgef-1::GFP + Prgef-1::sid-1],* CQ844: *set-2(ok952)* 2X outcrossed, IV183: *ueEx104 [odr-3::unc-31sense::sl2mCherry, odr-3::unc-31antisense::sl2mCherry, unc-122::RFP],* CQ757: *wqIs7 [Prgef-1::his-58::GFP],* PHX9664: *set-2(syb9664),* PY2417: *oyIs44 [odr-1::RFP],* PY3404: *oyIs48[ceh-36p::GFP],* CQ908 - *set-2 (ok952); wqEx96[Podr-1::SET1A+Pmyo-2::GFP],* CQ909 - *set-2(ok952); wqIs7 [Prgef-1::his-58::GFP],* CQ910 - *set-2(ok952); oyIs44 [odr-1::RFP],* MT6308: *eat-4(ky5)*.

### Auxin treatment

*set-2* was tagged on the C-terminus with a combined mIAA7 degron and GFP tag by CRISPR-Cas9-based genome editing. For auxin experiments, the standard HGM molten agar was supplemented with 2.5 mL 12 400 mM indole-3-acetic acid (IAA) per 1L of HGM: Alfa Aesar (#A10556) freshly prepared in ethanol and plates were seeded with E. coli for ad libitum feeding. Synchronized AID worms were transferred to HGM with auxin plates for 16-20 hours before behavior assay.

### RNAi treatment

RNAi experiments were performed using the standard feeding RNAi method. Bacterial clones expressing the control construct (empty vector, pL4440) and the dsRNA were obtained from the Ahringer RNAi library. All RNAi clones were sequenced prior to use.

### Pavlovian appetitive associative memory assay

Animals were trained and tested for short-term memory as previously described^25^. Briefly, synchronized young or aged adult hermaphrodites were washed from HGM, RNAi or HGM supplemented with auxin plates with M9 buffer, allowed to settle by gravity, and repeatedly washed three more times with M9 buffer. Then animals were starved for 1 hr in M9 buffer. For conditioning (food and 10% 2-butanone pairing), worms were then transferred to 10 cm NGM conditioning plates (seeded with OP50 *E. coli* bacteria and with 18 mL 10% 2-butanone (Acros Organics) dissolved in ethanol on the lid) for 1 hr at 20°C. After conditioning, the trained worms were tested for chemotaxis towards 10% butanone vs. an ethanol control either immediately (0 hr) or after being transferred to 10 cm NGM holding plates with fresh OP50 *E. coli* bacteria for specified time intervals.

Chemotaxis indices were calculated as follows:

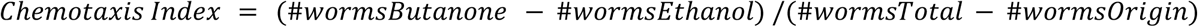

The calculation for the learning index is:

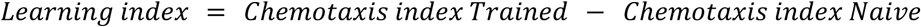

Learning indices for extrachromosomal transgenic strains were analyzed by hand counting GFP or mCherry positive and negative worms at different locations at individual timepoint on the chemotaxis plates. Control animals for these experiments were the transgenic worms’ GFP or mCherry negative siblings. For learning index bar plots, each dot indicates an individual chemotaxis assay plate containing ∼20-100 worms. 4-5 plates per time point per group were used.

To determine pharyngeal pumping rate (as a measure of appetite), wild type and *set-2 (ok952)* animals are starved and conditioned (as described), and the number of pharyngeal pumps is counted by visualization under a brightfield microscope after conditioning+2 hours hold on food.

### Chemotaxis Assay

Synchronized adult worms were tested for chemotaxis to 1% benzaldehyde in ethanol or 10% pyrazine in ethanol, using standard, previously described chemotaxis assay conditions^59^.

### Lifespan assay

Lifespan analyses were performed for whole-worm and neuron-specific *set-2* RNAi strains. RNAi-treated worms or their wild-type sibling controls were segregated at the L4 larval stage to start the assay. Every other day, worms were transferred to freshly seeded NGM plates with control or *set-2* RNAi bacteria. The first day of adulthood was defined at *T* = 0. Log-rank (Mantel–Cox) method was used to compare lifespans between transgenic and non-transgenic siblings. Worms that ‘escaped’ or ‘bagged’ were censored on the day of the event.

### Microscopy

For imaging of the *set-2::AID::GFP* strains, well-fed worms were transferred onto agar pads with sodium azide to visualize by Nikon AX-R microscope using the 488 nm laser. *z*-stack multi-channel (DIC, and GFP) images of Day 1 worms were acquired at ×60 magnification. Images were analyzed using Nikon NIS elements software.

### Actinomycin D treatment

For Actinomycin D experiments, worms were moved to NGM plates with either DMSO or 100 mg/mL Actinomycin D + DMSO after conditioning, following which they were tested for 1 and 2-hour memory.

### Optogenetics

Optogenetic activation of AWC was performed by adapting protocols described previously^36^.

L4-stage animals expressing Channelrhodopsin 2 in the AWC^37^ were incubated overnight on plates seeded with *E. coli* OP50 and 50 µM retinal (or *E. coli* and ethanol as control, as retinal is dissolved in ethanol). The following day, adult animals were conditioned with food and butanone. Following conditioning (0 hour), a subset of the population was held on plates with food for 2 hours (as for memory assays; Extended data Fig. 1) and another set was washed twice with M9. Individual animals were transferred onto unseeded NGM plates. Animals received 20-second pulses of blue light delivered by a Leica MZ16FA fluorescent compound microscope every two minutes, repeated eight times, and pulses five through eight were analyzed for the number of reversals. This was done at 0 and 2 hours after conditioning. Blinded analysis was performed with six animals per condition per replicate and a total of 3 replicates.

### WDR5-0103 treatment

For WDR5-0103 experiments, worms were transferred at the L4 larval stage onto HGM plates supplemented with 0.05M FUdR and 200uL of 10mM WDR5-0103 added on top of the bacterial lawn (leading to a final concentration of 100 mM). Following this, the worms were transferred every 48 hours to HGM+FUdR+ WDR5-0103 plates, and transferred back to standard HGM 20 hours before memory assay.

### AWC isolation by FACS

Wild type and *set-2* mutant young adult (Day 2) animals expressing *odr-1p::DsRed*^60^ (which brightly labels the AWC neurons with dimmer expression in AWB) were used to selectively sort the AWC neuron pair. This marker allowed us to sort a larger number of fluorescence-positive events by FACs (∼35,000 events/hour) at a higher rate (0.1% vs 0.05%) compared to a *ceh-36p::GFP*^61^ (labels AWC and ASE sensory neurons). The volumes and lyse times used for neuron isolation for fluorescence-based sorting were optimized to enable maximum number of fluorescence-positive events. Wild type and mutant animals were lysed at 0 and 2 hours after conditioning by adapting previously published protocols^39^. Briefly, worms were treated with 750 uL lysis buffer (200 mM DTT, 0.25% SDS, 20 mM HEPES pH 8.0, 3% sucrose) for 6.5 min to break the cuticle. Then worms were washed and resuspended in 450 uL 20 mg/mL pronase from *Streptomyces griseus* (Sigma-Aldrich). Worms were incubated at room temperature with mechanical disruption by pipetting until no whole-worm bodies were seen, and then ice-cold osmolarity-adjusted L-15 buffer (Gibco) with 2% Fetal Bovine Serum (Gibco) were added to stop the reaction. Prior to sorting, cell suspensions were filtered using a 5 um filter and DsRed+ cells were sorted using a BD Biosciences FACSAria Fusion sorter. Sorting gates were determined by comparing with age-matched, genotype-matched non-fluorescent cell suspension samples. Fluorescent neuron cells were directly sorted into Trizol LS.

### RNA extraction, library generation, and sequencing

We used the trizol-chloroform-isopropanol method to extract RNA, then performed RNA cleanup using RNeasy MinElute Cleanup Kit (Qiagen). RNA quality was assessed using the Agilent Bioanalyzer RNA Pico chip. 2 ng of RNA was used for library generation using Ovation SoLo RNA-Seq library preparation kit with AnyDeplete Probe Mix-*C. elegans* (Tecan Genomics) according to the manufacturer’s instructions (Barrett et al., 2021). Library quality and concentration was assessed using an Agilent Bioanalyzer DNA 12000 chip. Samples were multiplexed and sequencing was performed using NovaSeq S1 100nt Flowcell v1.5 (Illumina).

### ATAC-seq

*C. elegans* neuronal nuclei were isolated as previously^40^. FAC-sorted neuronal nuclei are pelleted by centrifuging at 1000g for 5 min at 4C. The supernatant was removed without disrupting the pellet. 100uL of chilled lysis buffer (10 mM Tris-HCl (pH 7.4), 10 mM NaCl, 3 mM MgCl_2_, 0.1 % Tween-20, 0.1 % NP-40 substitute, 0.01 % Digitonin, 1 % BSA, nuclease-free H_2_O to 2 mL) was added, mixed by pipetting up and down 10 times, and incubated for 1 min on ice. After incubation, 1 mL chilled Wash Buffer (10 mM Tris-HCl (pH 7.4), 10 mM NaCl, 3 mM MgCl_2_, 1% BSA, 0.1% Tween-20 in nuclease-free H_2_O) was added to the lysed cells, mixed by pipetting up and down 5 times, and centrifuged at 1000g for 10 min at 4C. The supernatant was removed without disrupting the pellet. 50uL ATAC Transposition reaction was added to the lysed pellet, and incubated at 37C for 25 min. Following transposition, the tagmented fragments were purified using the Qiagen MinElute PCR purification kit. Sequencing was performed using 101-bp paired-end sequencing on an Illumina Novaseq S1 100nt flowcell v1.5 platform. The ATAC-seq libraries were sequenced to a median depth of over 40 million unique, high-quality mapping reads per sample. Prior to mapping, standard next-generation sequencing quality control steps, as well as ATAC-seq-specific quality control steps, were performed.

### Alignment with human WDR5 and SET1A

The crystal structure of human WDR5 (PDB: 3P4F) and the predicted AlphaFold structure of C. elegans WDR-5.1 (UniProt ID: P61964, shown in pink) were loaded into PyMOL (Schrödinger, LLC) and aligned using the *align* and *cealign* algorithms, yielding an RMSD of 0.57 Å across 304 residues. To assess conservation of the ligand-binding site, the crystal structure of WDR5 complexed with WDR5-0103 (PDB: 3UR4, shown in blue) was loaded into PyMOL and overlaid with C. elegans WDR-5.1. The resulting superposition suggests that the ligand-binding pocket is structurally conserved and that WDR5-0103 may interact with C. elegans WDR-5.1 in a manner similar to that observed for human WDR5.

Full-length AlphaFold structures of human SET1A (AF-O15047-F1-v6) and *C. elegans* SET-2 (AF-Q18221-F1-v6) were also loaded into PyMOL and aligned using the *align* and *cealign* algorithms. Alignment of the full-length proteins yielded an RMSD of 4.70 Å across 224 residues. In contrast, alignment restricted to the catalytic domains produced an RMSD of 0.48 Å across 136 residues, indicating that the catalytic domains of SET1A and SET-2 occupy nearly identical positions in three-dimensional space. This high structural similarity suggests that the catalytic architecture is strongly conserved between the two proteins.

### RNA-seq data analysis

RNA sequencing analysis was performed as previously described. FASTQC was used to assess read quality scores. The universal Illumina adaptor sequences were trimmed using Cutadapt v1.6. The trimmed reads were mapped to the *C. elegans* genome (UCSC Feb 2013, ce11/ws245) using STAR. The reads aligning to individual genes were counted using htseq-counts (mode: union), and DESeq2 was used for differential expression analysis. Genes with an adjusted *p-value* ⋜ 0.05 were considered significantly differentially expressed.

### Gene Ontology Analysis

Functional categories for the genes expressed in wild type AWC at 0 hours were determined using WormCAT^54^. Genes with log(Mean DeSeq2 Normalized Counts)>2 were defined as expressed. gProfiler^57^ was used for ontology (GO) term analysis from upregulated or downregulated gene lists (DESeq2 adjusted *p-value* ⋜ 0.05).

### Tissue query

worm.princeton.edu^43^ was used for tissue query from upregulated or downregulated gene lists (DESeq2 adjusted *p-value* ⋜ 0.05).

### ATAC-seq data analysis

Prior to mapping, standard next-generation sequencing quality control steps, as well as ATAC-seq-specific quality control steps, were performed. ATAC-seq reads were aligned using Bowtie2 and peaks were called using MACS3. Peak margins were defined and differential read counts determined using bedtools. Differential accessibility analysis was performed using DeSeq2 in R. Differentially accessible regions (DARs) were determined by FDR-adjusted p-values<0.05. Peak-to-gene mapping was performed using the bedtools closest function. PCA plots generated in R and volcano plots were generated using pandas. The code used for data analysis was adapted from https://github.com/guertinlab^62^.

### Statistical analysis

For all comparisons between genotypes and across timepoints, two-way ANOVA was conducted. If interaction between genotype and time was significant (p<0.05), simple main effects analyses were done, comparing chemotaxis indices of all the genotypes at all time points. When 2 genotypes are present, there is only one pairwise comparison at each time point, and Šidák correction was used for multiple comparisons, with adjusted p-values reported. When >2 genotypes are present, there are multiple pairwise comparisons at each time point, and Tukey’s post hoc tests were done, and adjusted p-values reported. For memory time-courses, the adjusted p-values corresponding to the 1- or 2-hr timepoints are reported on the 1- or 2-hr learning index bar plots. Experiments were repeated on separate days, using separate independent populations, to confirm that results were reproducible. Prism 9 software was used for all statistical analyses. DeSeq2 was used for differential expression (RNA-seq) and differential accessibility (ATAC-seq) analysis. For lifespan assays, Log-rank (Mantel-Cox) test was used.

To determine the required sample size for comparing chemotaxis indices between genotypes and across time points for memory assays, we conducted a power analysis for a repeated-measures ANOVA (two-sided a=0.05). Based on a typical mean difference=0.2 and standard error=0.05^38^, 5 technical replicates per time point per group is required for 80% power; therefore, we used 5 assay plates (150-200 worms/plate) per time point. For pharyngeal pumping assays, 4 samples per group are required for 80% power at a=0.05^63^ for an expected difference in means=50 pumps/min and standard error=10 pumps/min (using unpaired t-test); 8 worms per group were used, ensuring sufficient power. In lifespan assays, for an expected median lifespan difference=20%, a sample size=100 per group is required for 80% power at a=0.05 (using log rank test)^21^; 120 worms were used per group, ensuring sufficient power. Sample sizes are consistent with previous studies.

## Data availability

The manuscript contains all the data required to evaluate its conclusions. RNA-seq (PRJNA1436608) and ATAC-seq (PRJNA1436611) data have been deposited in NCBI.

