## Supplementary material for "COMPASS-mediated epigenetic regulation of transcription accelerates forgetting with age": Supp Figs 1-6

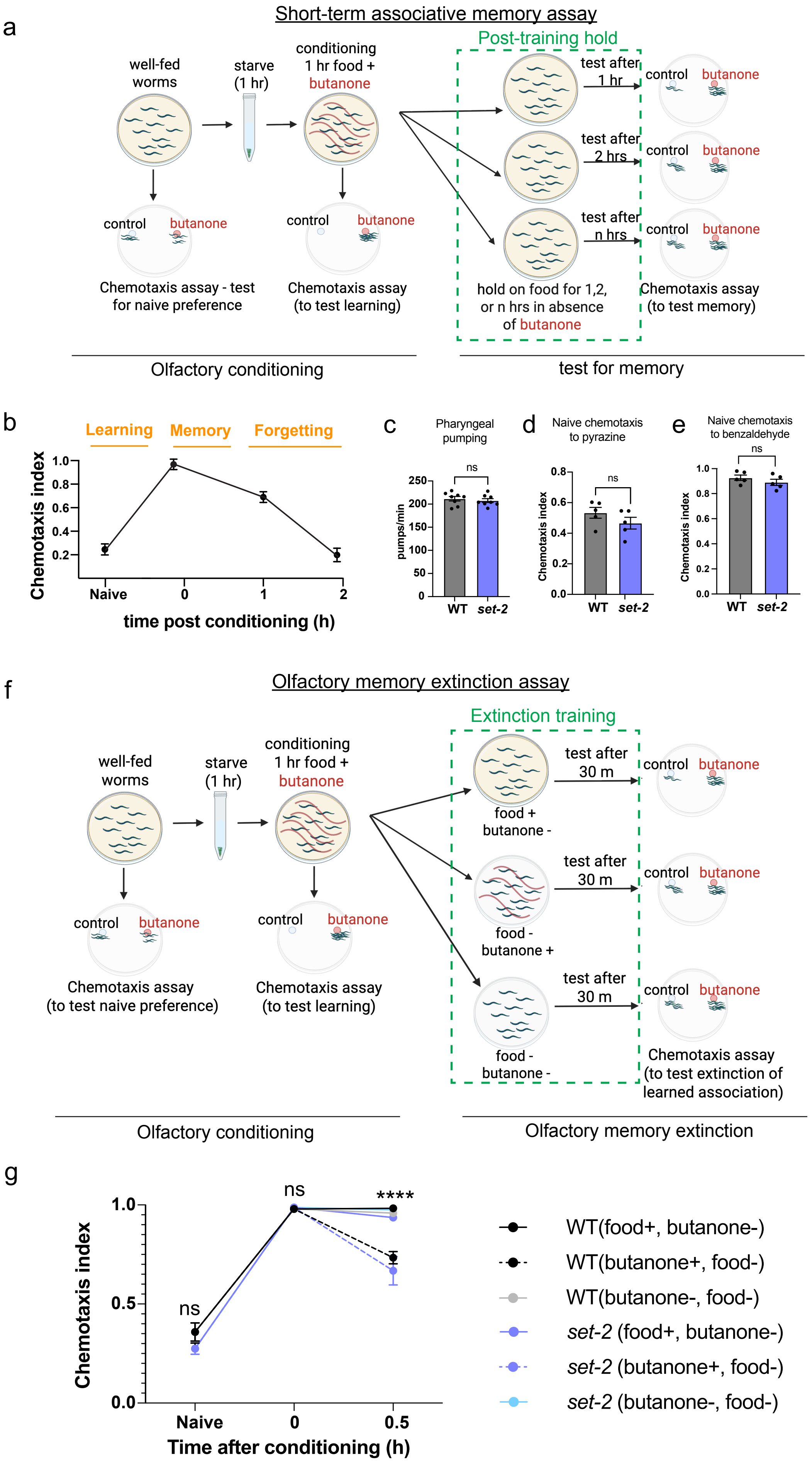


Extended data Figure 1

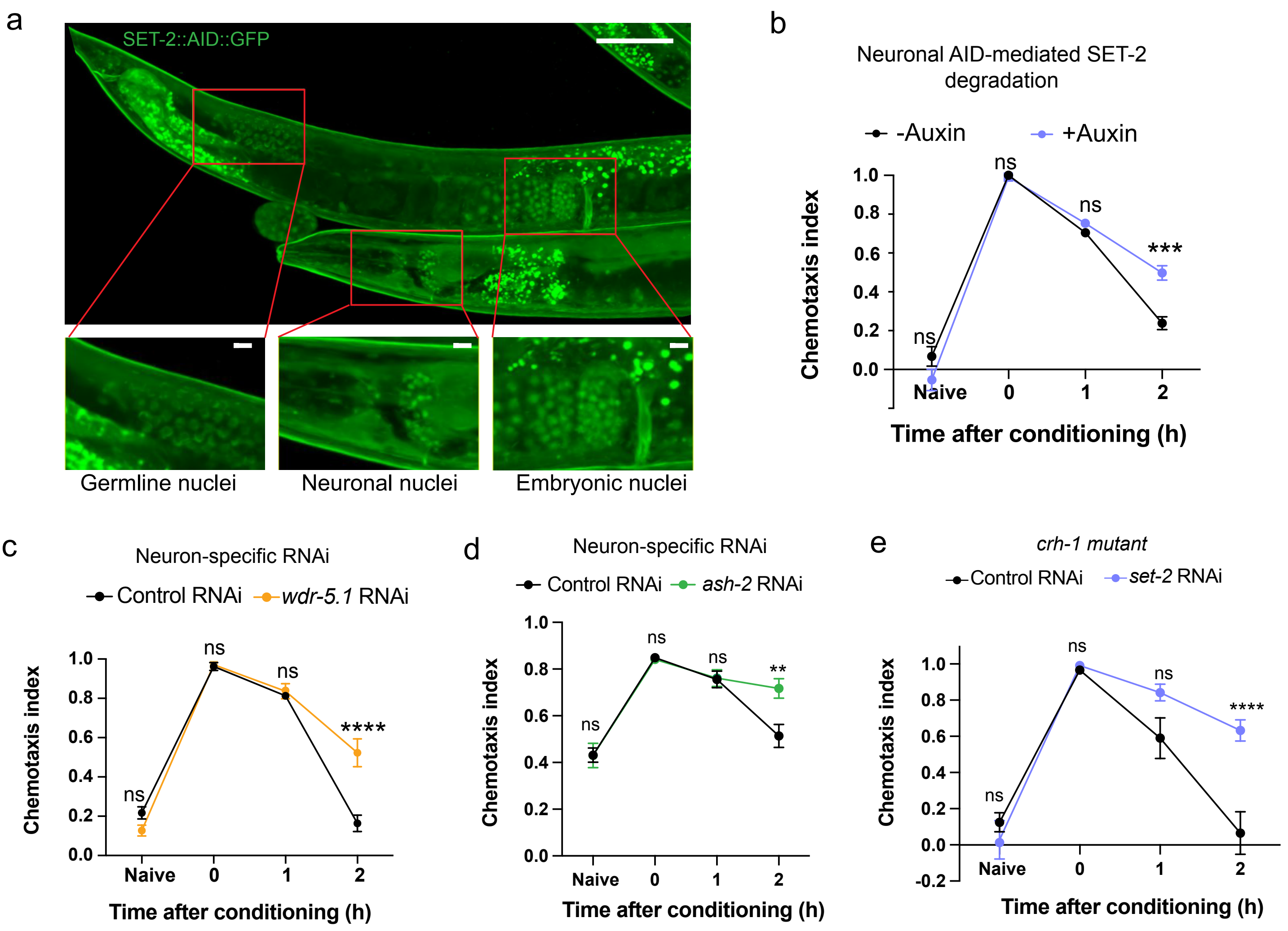


Extended data Figure 2

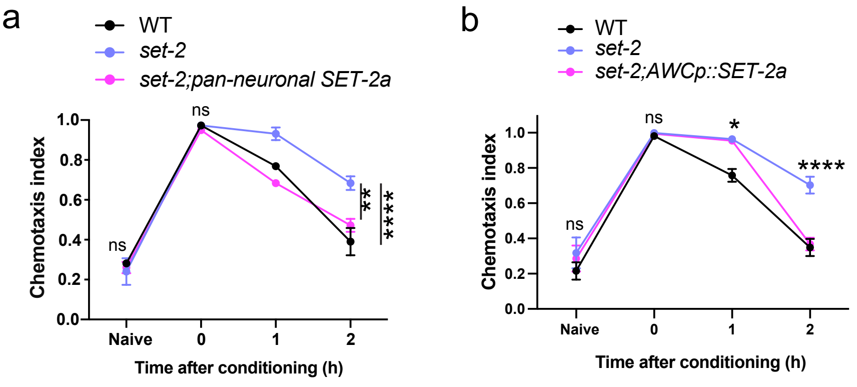


c

AWC-expressed genes ranked by normalized counts

| Gene | Mean Normalized Counts | Function |
|---|---|---|
| eef-1A.1 | 226094.5613 | Translation elongation |
| unc-44 | 199504.4018 | Cell adhesion molecule binding |
| ctc-1 | 175918.8498 | Heme and metal ion binding |
| srpr-2.2 | 167390.892 | Unknown; neuron-enriched |
| MTCE.33 | 165949.6674 | Mitochondrial 23Ss ribosomal RNA |
| srpr-2.1 | 161851.2746 | Unknown; neuron-enriched |
| egl-3 | 136189.8458 | Neuropeptide processing |
| ZK185.9 | 109918.7598 | Unknown |
| hsp-1 | 108175.6846 | ATP hydrolysis activity; heat shock response |
| ctc-3 | 107229.7302 | Electron transfer; cytochrome c oxidase activity |
| unc-54 | 105099.3339 | Muscle myosin class II heavy chain |
| lfi-1 | 101971.5742 | Located in kinetochore microtubule and nucleus |
| act-4 | 90482.22646 | Actin isoform |
| eel-1 | 88804.51758 | Ubiquitin protein ligase activity |
| eef-2 | 82643.46683 | Translation elongation |
| cbd-1 | 80506.64395 | Chitin binding |
| cpg-1 | 72483.76275 | Chitin binding |
| casy-1 | 71941.16556 | Calcium ion binding |
| hsp-90 | 70664.85647 | Molecular chaperone |
| cab-1 | 67815.88336 | Chemical synaptic transmission |
| apl-1 | 66737.5005 | Orthologous to human Amyloid Precursor Protein |
| egl-21 | 65982.04486 | Neuropeptide processing |
| cpg-2 | 65594.02253 | Chitin binding |
| pab-1 | 65312.10255 | mRNA binding |
| tbb-2 | 62149.5934 | Constituent of microtubule cytoskeleton |
| nlp-3 | 61115.10469 | Neuropeptide signaling |
| epic-2 | 60439.46658 | Unknown |
| ant-1.1 | 60112.28187 | Mitochondrial adenine nucleotide transporter |
| nrsn-1 | 59446.48773 | Nervous system development |
| ida-1 | 59088.81036 | Constituent of dense-core vesicle membrane |
| sbt-1 | 57484.19583 | Peptidase regulation |
| unc-70 | 56696.57285 | Beta-spectrin ortholog |
| vig-1 | 55568.43991 | RNA-binding |
| atp-1 | 53773.0029 | Constituent of mitochondrial ATP synthase |
| pgal-1 | 52014.20478 | Peptide metabolic process |
| ctb-1 | 52014.20478 | Mitochondrial complex III subunit |
| atp-2 | 48396.48527 | ATP synthase subunit |
| snt-4 | 48046.68576 | SNARE-binding activity |
| gpb-1 | 46514.10369 | G-protein beta subunit; mitotic spindle regulation |
| R02F2.1 | 46001.23727 | Cyclin-dependent protein serine/threonine kinase |

mitochondrial constituent
Neuropeptide processing
Cytoskeletal function
Chitin binding
Neuronal function: General
heat shock response
Translation elongation
RNA binding
Synaptic function

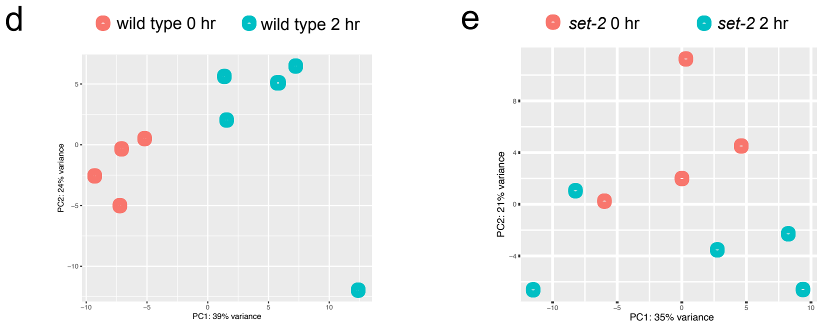


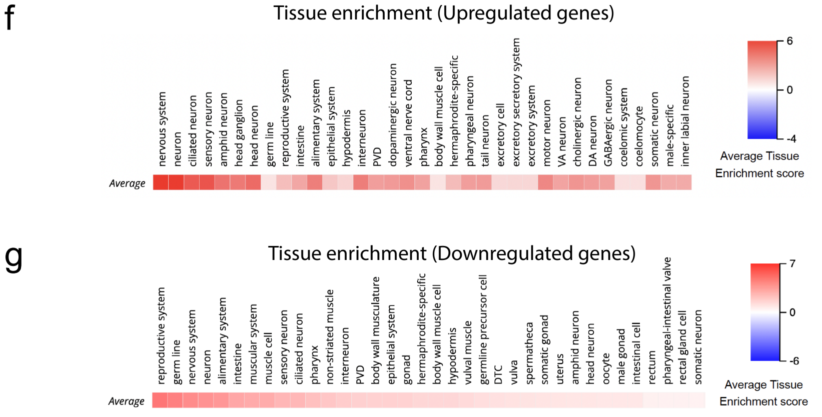


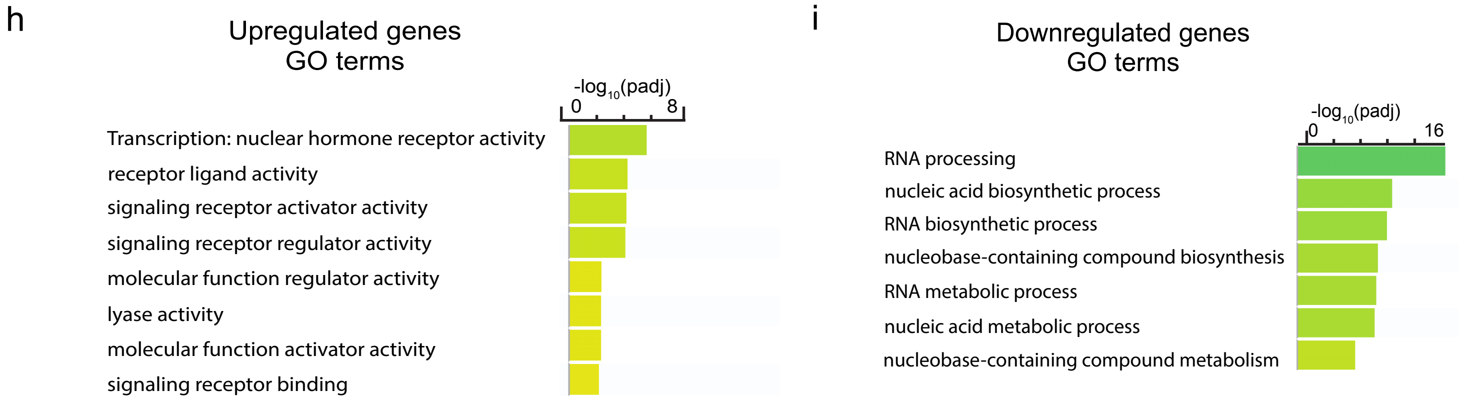


### Extended data Figure 3

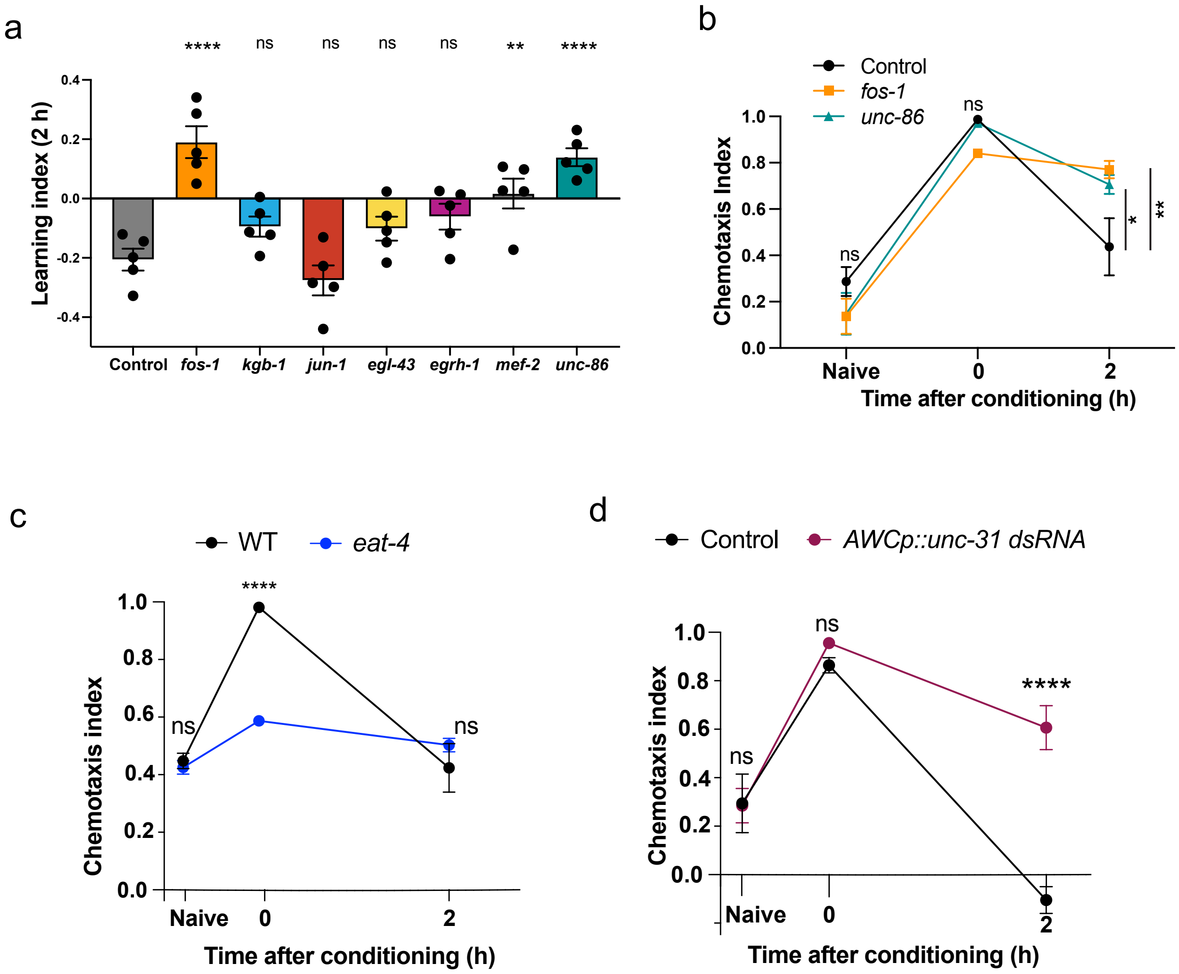


Extended data Figure 4

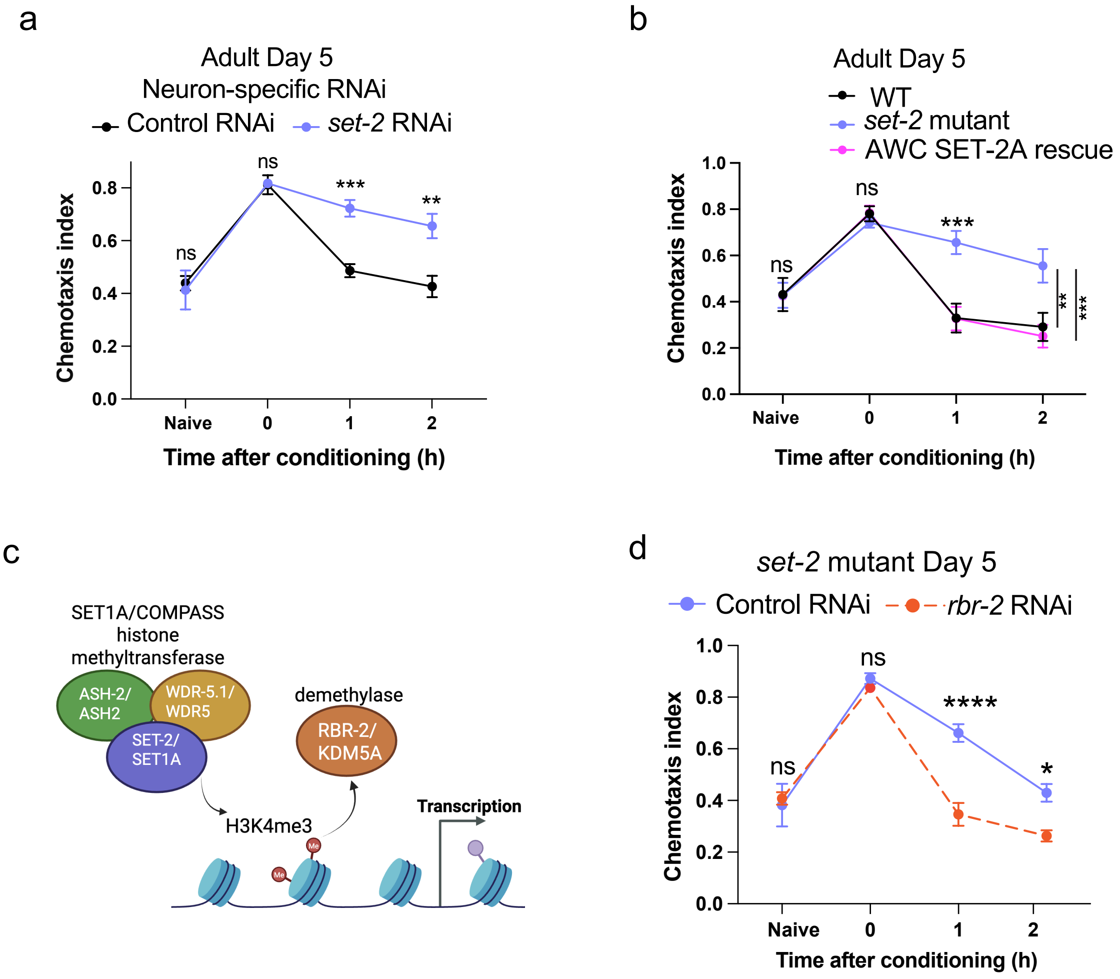


Extended data Figure 5

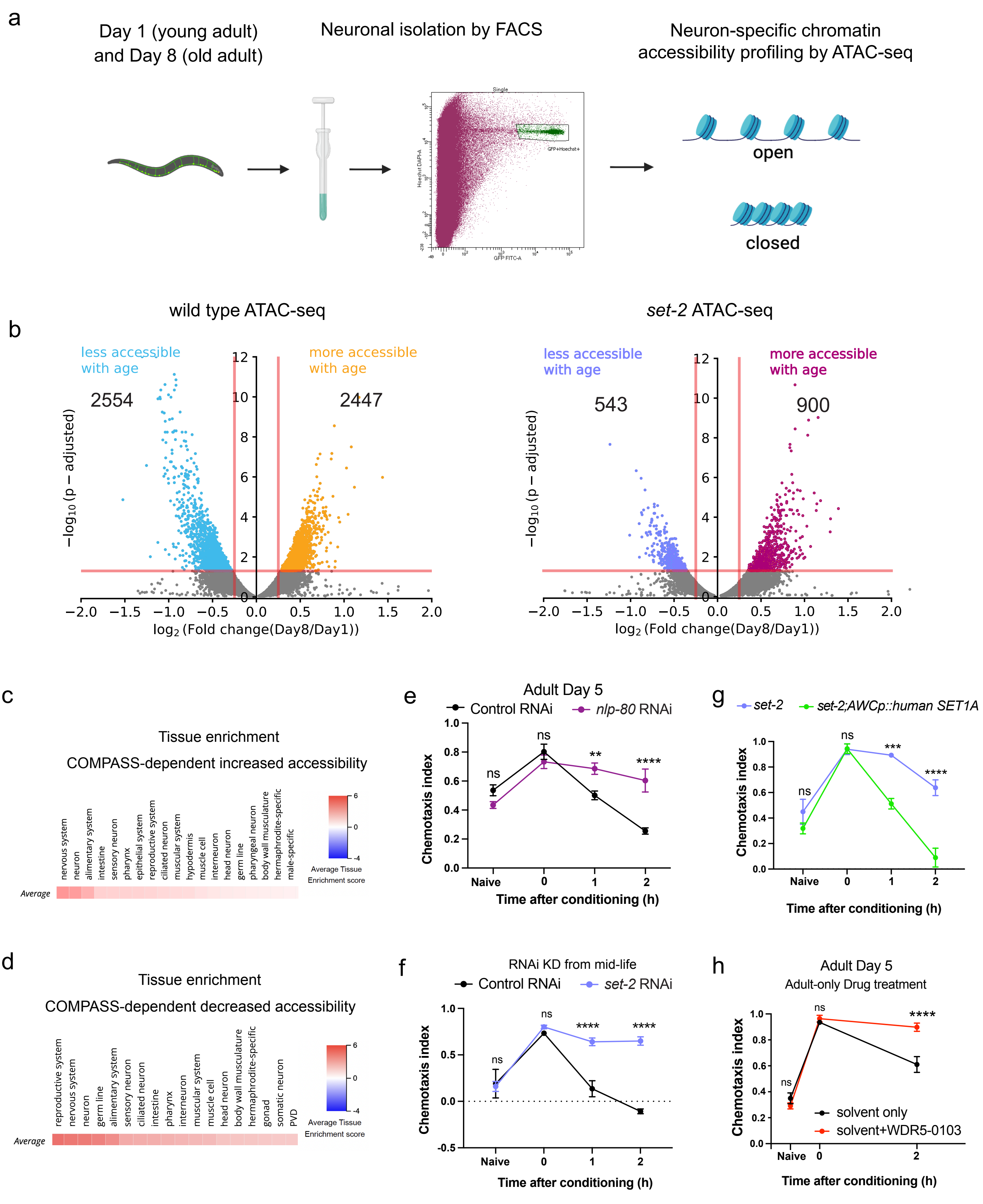


### Extended data Figure 6
