## Supplementary material for "COMPASS-mediated epigenetic regulation of transcription accelerates forgetting with age": File S1

### BD FACSDiva 9.0.1

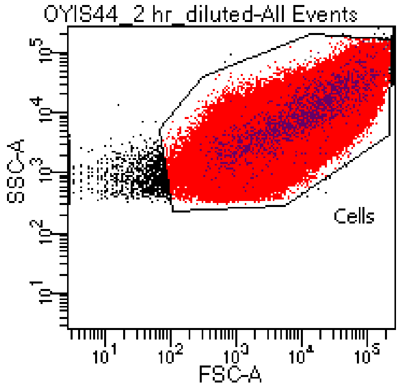


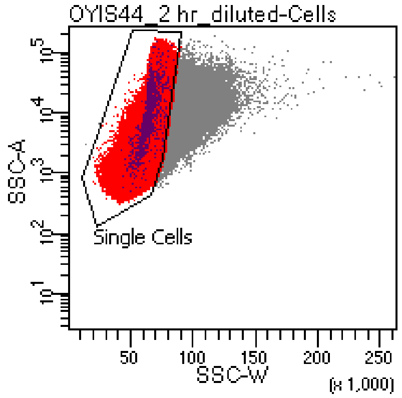


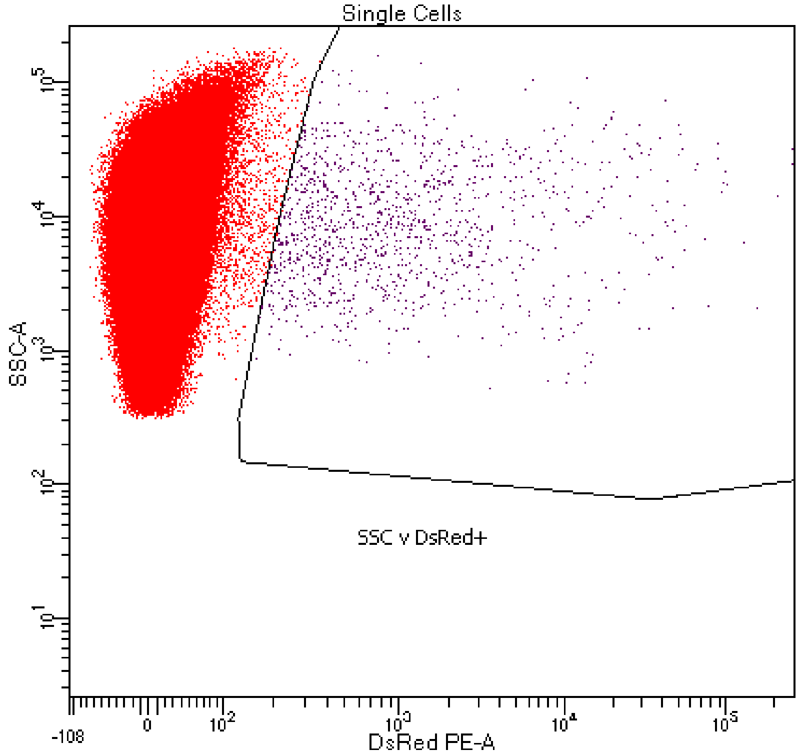


Tube: OYIS44_2 hr_diluted

| Population | #Events | %Parent | %Total |
|---|---|---|---|
| All Events | 1,033,729 | #### | 100.0 |
| Cells | 1,031,479 | 99.8 | 99.8 |
| Single Cells | 1,000,000 | 96.9 | 96.7 |
| SSC v DsRed+ | 1,232 | 0.1 | 0.1 |

Experiment Name: TS_20240930_ODR WT_DsRed_ neurons
Tube Name: OYIS44_2 hr_diluted
Record Date: Sep 30, 2024 5:15:00 PM

| Population | #Events | %Parent | GFP FITC-A Median | DsRed PE-A Median |
|---|---|---|---|---|
| Single Cells | 1,000,000 | 96.9482 | 4 | 5 |
| SSC v DsRed+ | 1,232 | 0.1232 | 14 | 807 |
