## Supplementary material for "COMPASS-mediated epigenetic regulation of transcription accelerates forgetting with age": File S2

### BD FACSDiva 9.0.1

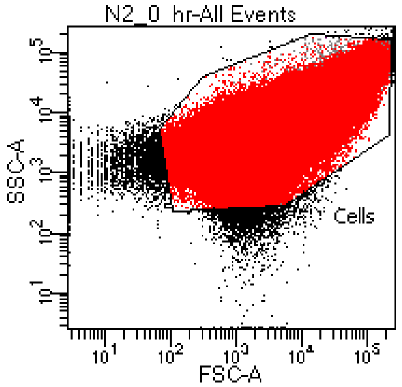


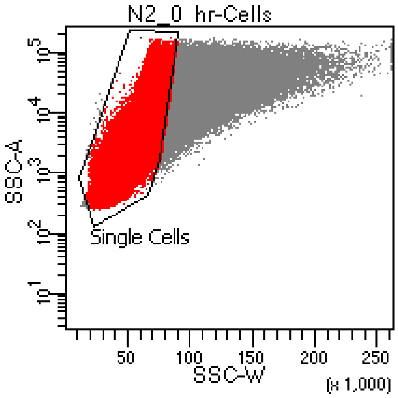


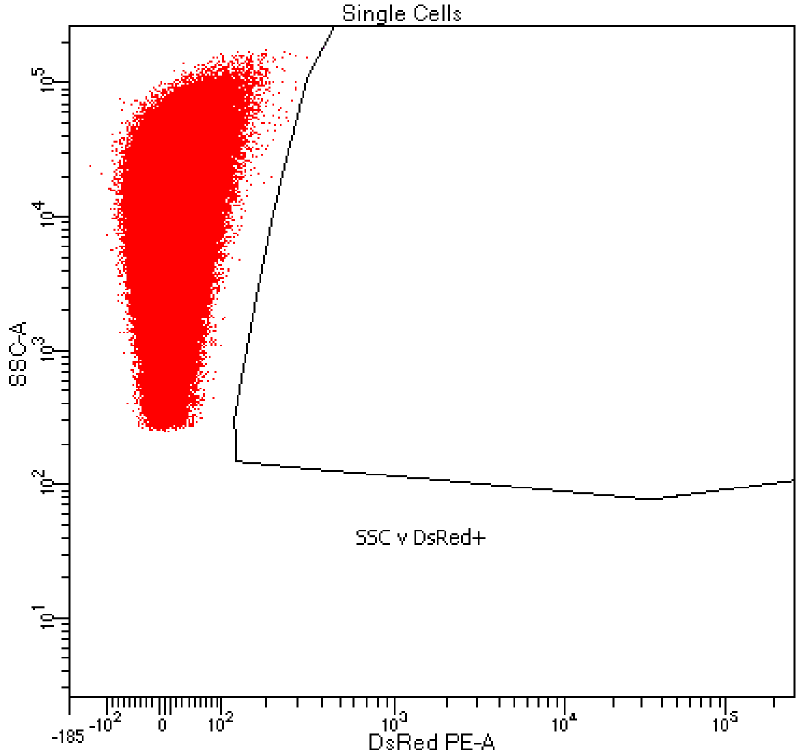


Tube: N2_0 hr

| Population | #Events | %Parent | %Total |
|---|---|---|---|
| All Events | 1,225,257 | #### | 100.0 |
| Cells | 1,211,749 | 98.9 | 98.9 |
| Single Cells | 1,000,000 | 82.5 | 81.6 |
| SSC v DsRed+ | 1 | 0.0 | 0.0 |

Experiment Name: TS_20240930_ODR WT_DsRed_ neurons
Tube Name: N2_0 hr
Record Date: Sep 30, 2024 3:02:39 PM

| Population | #Events | %Parent | GFP FITC-A Median | DsRed PE-A Median |
|---|---|---|---|---|
| Single Cells | 1,000,000 | 82.5253 | 5 | 7 |
| SSC v DsRed+ | 1 | 0.0001 | -10 | 401 |
